# Preponderance of generalized chain functions in reconstructed Boolean models of biological networks

**DOI:** 10.1101/2023.10.08.561412

**Authors:** Suchetana Mitra, Priyotosh Sil, Ajay Subbaroyan, Olivier C. Martin, Areejit Samal

## Abstract

Boolean networks (BNs) have been extensively used to model the dynamics of gene regulatory networks (GRNs) that underlie cellular decisions. The dynamics of BNs depend on the network architecture and *regulatory logic rules* (or *Boolean functions* (BFs)) associated with nodes, both of which have been shown to be far from random in large-scale studies of reconstructed Boolean models. At the level of the BFs, nested canalyzing functions (NCFs) have been shown to be strongly enriched in such GRN models. The central question we address here is whether that enrichment is due to certain sub-types of NCFs. To answer this, we build on one sub-type of NCFs, the *chain functions* (or *chain-0 functions*) proposed by Gat-Viks and Shamir. First, we propose 2 other sub-types of NCFs, namely, the class of *chain-1 functions* which is the dual of the class of chain-0 functions, and *generalized chain functions*, the union of the chain-0 and chain-1 types. Next, we find that the fraction of NCFs that are chain-0 functions decays exponentially with the number of inputs, and exhibits a fractal-like behaviour as a function of the bias for a fixed number of inputs. Moreover, we explain several of these observations analytically. Then, by analyzing 5990 BFs extracted from a large dataset of reconstructed Boolean models, and 2 other datasets, we find that generalized chain functions are significantly enriched within the NCFs. Lastly, we illustrate the severe restriction imposed by generalized chain functions compared to NCFs for 3 biological models and perform model selection on them using known relative stability constraints.

## 1. INTRODUCTION

Cellular decision making processes are governed by intricate gene regulatory networks (GRNs) [1]. Extensive efforts have been dedicated in the pursuit of understanding their underlying structure and dynamics [2–7]. The discrete state framework of “logical modeling” pioneered by Stuart Kauffman [8, 9] and René Thomas [10, 11], stands out as a simple and effective way to mimic the dynamic behaviour of GRN. In the Boolean formalism, the state of the nodes (genes or other biological entities) are simplified to two different states -’off’ or ‘on’.

In order to replicate key steady-state dynamical behaviour in living systems such as fixed points, Kauffman explored Random Boolean networks (RBNs) [8]. RBNs are defined by the inclusion of interactions (directed edges) between nodes (genes) chosen at random, accompanied by the assignment of random logical update rules at these nodes. However, extensive studies of biological networks, aided by recent advances in collection of large-scale data, have revealed that these network architectures are far from being random, both in terms of their topology and the logic rules [4, 6, 7, 12, 13]. Since the beginning of the present century, there has been a significant surge in employing the Boolean framework to reconstruct GRNs from experimental biological data [14–17]. This trend has been propelled by the advancement in sequencing technology and increased computational capabilities, allowing not only the reconstruction of networks but also in reproduction of gene expression patterns.

Deeper investigations into the use of BFs has unveiled that specific classes of functions, such as unate functions (UFs) [18], canalyzing functions (CFs) [2] and NCFs [3, 19] exhibit distinct properties that inherently render them more suitable to be regarded as biologically meaningful when compared to random BFs. Notably, some of us in our previous work have shown that two classes of BFs, namely NCFs and read-once-functions (RoFs) exhibit unexpected prevalence [13] within a compiled reference biological dataset of 2687 BFs derived from reconstructed Boolean models of biological systems despite their theoretical fraction within the space of all BFs being minuscule. We also showed that NCFs and RoFs have the ‘simplest logic’ with respect to the complexity measures ‘average sensitivity’ [20] and ‘Boolean complexity’ [21] respectively. In a more recent work, we have shown that such biologically meaningful logics lead to more bushy and convergent dynamics, indicative of a higher degree of order [22].

In this study, our focus centers on two specific sub-types of NCFs. One is known as the chain function class that was introduced by Gat-Viks and Shamir in 2003 [23]. They argued for the ubiquity of these functions in biological networks. Furthermore, in the same year, Kauffman *et al*. [3] provided a simplified definition of the chain function class based on the canalyzing input values. Their work revealed that out of the 139 rules (or BFs) compiled by Harris *et al*. [24], 132 were NCFs and amongst those 107 were chain functions. In 2011, Akutsu *et al*. [25] proposed an alternative definition of chain functions based on the Boolean expression, where the variable signs and the subsequent operators are strictly interdependent and governed by a control pattern that is a sequence of ‘0’ and ‘1’. In our current work, we present an equivalent definition of Akutsu *et al*. [25], where we use the bias of a Boolean function instead of their control pattern. We also introduce a novel sub-type of NCFs, comprised of the duals of the functions in the class of chain functions. To distinguish the class of chain functions from this new class, we refer to the former as chain-0 functions or *ChF*_0_, and the latter one as chain-1 functions or *ChF*_1_. We then term the union of these two classes as ‘generalized chain functions’ and denote it by *ChF*_*U*_. We first derive a formula to count the number of functions of a *k*-input *ChF*_0_ (or *ChF*_1_) with bias *P*. We similarly investigate the fraction of NCFs that belong to *ChF*_0_ and *ChF*_1_. Next we assess the presence of functions from these two classes in more extensive and contemporary datasets derived from reconstructed biological Boolean networks (BNs). For this analysis, we utilize three reference biological datasets: (a) BBM benchmark dataset [26] which is the most recent and largest repository of regulatory logic rules from which we extract 5990 BFs, (b) MCBF dataset [13], comprising of 2687 BFs, compiled previously by some of us from 88 published biological models, and (c) Harris dataset [24], comprising of 139 BFs. Furthermore, we demonstrate the practical utility of these special sub-types of NCFs in the context of model selection. In prior work [27], we have shown that confining BFs to NCFs can yield a substantial reduction in the space of plausible models. However, in that work we showed that even for networks with 18 nodes, the search space remained sufficiently vast (even when restricting to NCFs), making exhaustive search infeasible. Therefore we ask whether the generalized chain functions can further curtail the space of potential candidate models. To do so, we perform a comprehensive case study using three biological models [28–30], and finally use relative stability as a constraint to select for models within ensembles that employ chain functions.

## 2. METHODS

### 2.1. Boolean models of Gene Regulatory Networks

A Boolean model of a GRN consists of nodes and directed edges where nodes correspond to genes and directed edges correspond to the regulatory interactions between them [2, 8, 9]. Genes in a BN are either in an upregulated (‘on’) or downregulated (‘off’) state. In a BN with *N* nodes, we denote the state of the *i*^*th*^ gene at time *t* by *x*_*i*_(*t*), where *i* ∈ {1, 2, …, *N*} and *x*_*i*_(*t*) ∈ 0, 1 . The state of the network at time *t* can be given by a vector **X**(*t*) = (*x*_1_(*t*), *x*_2_(*t*), …, *x*_*N*_ (*t*)). The temporal dynamics of a BN are dictated by the BFs (or *logical update rules* or *regulatory logic*) along with an update scheme (*synchronous* [2] or *asynchronous* [11]). In the synchronous update scheme, which is the mode of update used in this work, all nodes of the BN are updated simultaneously. Mathematically, this may be expressed as 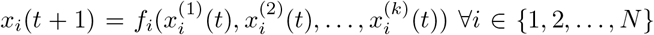, where *f*_*i*_ is the BF that acts on the *k* inputs to node *i, j* ∈ 1, {2, …, *k*} and 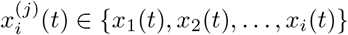. This type of local dynamics takes the network from the state **X**(*t*) to the state **X**(*t* + 1). Such a scheme leads to two kinds of emergent dynamics. In the first, the system reaches a state which on the next (and subsequent) update is left unchanged, corresponding to a *fixed point attractor*. In the second, the system cycles infinitely through a fixed set of states on successive updates, corresponding to a *cyclic attractor*. The states that converge to an attractor (including the attractor itself) comprise its basin of attraction. Of these two types of attractors, fixed points are considered the most relevant ones from a biological point of view; in multi-cellular organisms, one considers that they provide the gene expression patterns characteristic of different cellular types [2].

### 2.2. Representations and properties of Boolean Functions

#### 2.2.1. Truth Table and Output Vector

A BF *f* with *k* inputs (which we also call as a *k*-input BF) can be specified via a truth table with *k* + 1 columns and 2^*k*^ rows, where the first *k* columns correspond to the states of the input variables and the (*k* + 1)^*th*^ column corresponds to the output values. Each row of the truth table is a unique combination of the states of *k* input variables with the (*k* + 1)^*th*^ entry corresponding to the output value associated with that input combination. The output column of the BF can also be considered as a binary vector with 2^*k*^ elements. The bias *P* of a BF is the number of 1*s* in its binary output vector. It is sometimes convenient to specify that output vector via the integer whose binary decomposition is given by the entries of that vector. Note that for a *k*-input BF there are 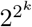 distinct BFs possible since the vector has 2^*k*^ entries.

#### 2.2.2. Boolean Function Expression

A *k*-input BF can also be represented as a Boolean expression that combines Boolean variables (*x*_*i*_ ∈ {0, 1}) with the logical operators conjunction (or AND or ∧), disjunction (or OR or ∨) and negation (or NOT or 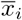). For illustration 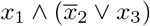 and 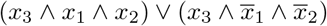 are cases of 3-input BFs.

#### 2.2.3. Operations on Boolean functions

##### Negation of variables

Given a *k*-input BF *f*, one can consider the modified function where some of the input variables are negated. There are 2^*k*^ possible negation operations if we include the negation of none of the inputs. Such a negation may or may not lead to a new BF but preserves the bias of the BF. For example if 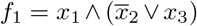 and 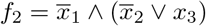, then one function can be obtained from the other by negating the variable *x*_1_.

##### Permutation

Given a *k*-input BF *f*, a permutation operation is performed by permuting the variables in the BF’s logical expression. There are *k*! possible permutation operations when one includes the identity permutation. Such an operation may or may not lead to a new BF but it preserves the bias of the BF. For example if 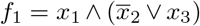 and 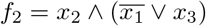, then one function can be obtained from the other by permuting the variables *x*_1_ and *x*_2_.

##### Complementation

Given a *k*-input BF *f*, a complementation operation replaces the 0*s* and 1*s* of the output column of the truth table with 1*s* and 0*s* respectively. In terms of the Boolean expression, this is equivalent to negating all the variables and changing the ∧ and ∨ operators to ∨ and ∧ operators respectively. For example, the BFs 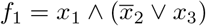 and 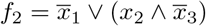 are complements of each other.

### 2.3. Nested Canalyzing Functions

#### Definition 1 (NCF)

A *k*-input BF is *nested canalyzing* [3, 31] *with respect to a permutation σ* on its inputs {1, 2, …, *k*} if:

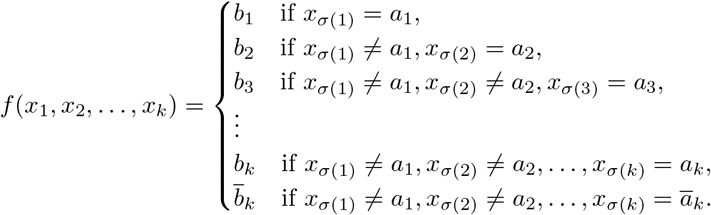

In the above equation, *a*_1_, *a*_2_, …, *a*_*k*_ are the canalyzing input values and *b*_1_, *b*_2_, …, *b*_*k*_ are the canalyzed output values for inputs *σ*(1), *σ*(2), …, *σ*(*k*) in the permutation *σ* of the *k* inputs. Here, *a*_*k*_ and *b*_*k*_ are the complements of the Boolean values *ā*_*k*_ and 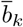, respectively.

Alternatively, NCFs can be represented succinctly via the following Boolean expression [32]:

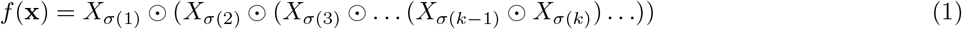

where 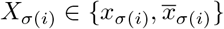 and ⊙ ∈ {∧, ∨}.

The Boolean expression in Eq. (1) may also written to regroup the successive OR and AND logical operators:

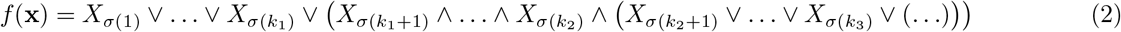

or, if the first logical operator is an AND:

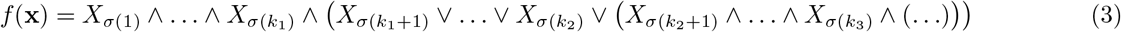

#### Definition 2 (Layer)

Given a *k*-input NCF *f* with respect to a permutation *σ* on its inputs {1, 2, …, *k* }, we can make explicit the consecutive canalyzing output values as follows:

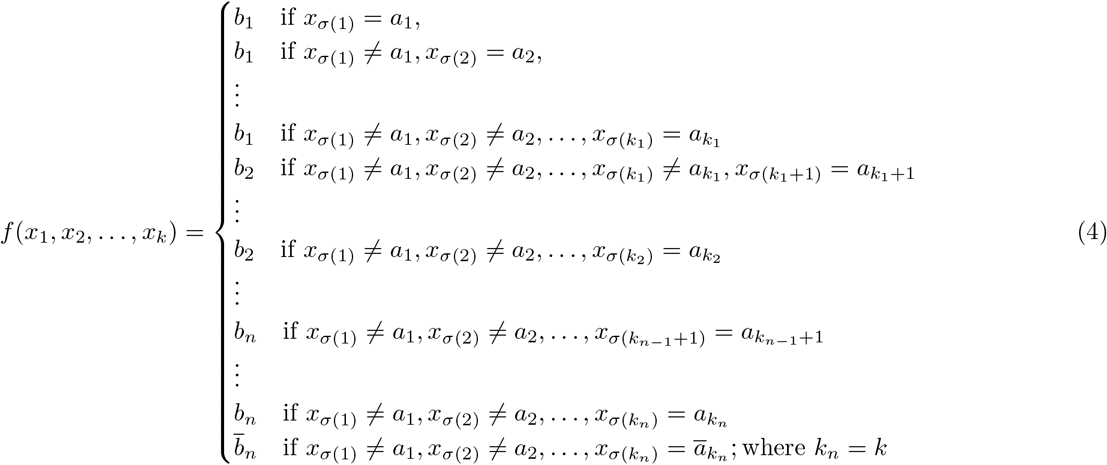

where, 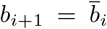 for all *i* ∈ {1, 2, …, *n*} . The consecutive canalyzing inputs giving the same canalyzed output are grouped into what is referred to as a **layer** [33]. The number of inputs present in a layer can be termed the *layer-size* and the number of layers called the *layer-number*. Hereafter, we will denote the layer-size of the *i*^*th*^ layer as *m*_*i*_, the layer-size of the last layer being *m*_*last*_. In the notation of Eq. (4), *m*_1_ = *k*_1_, *m*_2_ = *k*_2_−*k*_1_, …, *m*_*last*_ = *k*_*n*_−*k*_*n*−1_.

An equivalent definition of the layers can be provided based on the logical expression of a NCF. Specifically, each successive set of inputs followed by the same type of operators (‘∧’ or ‘∨’) constitutes a distinct layer (see Eqs. (2), (3)). In other words, in the expression of a NCF, whenever the operator flips from AND (∧) to OR (∨) or vice-versa, a new layer begins (including the preceding variable of the flipped operator). For example, given the NCF 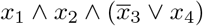, we immediately observe that it has two layers with *m*_1_ = *m*_2_ = 2.

#### 2.3.1. Relationship between bias, operators and layers

We now formalize the relationship between the bias and the sequence of operators in the Boolean expression (Eq. (1)) for NCFs. In [32], the authors encoded the sequence of operators in the Boolean expression of *k*-input NCF (Eq. (1)) with *k* − 1 bits, but did not relate their encoding to the bias *P* of the NCF. This relationship between the bias and operator sequence was implicitly first used in [13] but was not stated explicitly. The bias *P* of a *k*-input NCF can be expressed via its binary representation with *k* bits where the least significant bit is 1 since NCFs have odd bias. The first (*k*−1) bits of this binary string encode the operator sequence that appears in the associated Boolean expression of NCFs (Eq. (1)) such that the bits 0 and 1 encode the operators ‘∧’ and ‘∨’ respectively. We explain the relationship in more detail in Appendix A, Property A.1. Let us illustrate this relation via an example. Consider a 4-input NCF with bias *P* = 5. The associated binary string representation of *P* is 0101. The Boolean expression for a 4-input NCF with *P* = 5 is *X*_*σ*(1)_ ∧ (*X*_*σ*(2)_ ∨ (*X*_*σ*(3)_ ∧ *X*_*σ*(4)_)) where 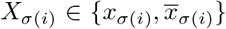. The operator sequence for this expression is (∧,∨,∧) (starting from the outermost operator to the innermost one), while the initial (*k*−1) bits of the binary representation are 010.

For any NCF, both the layer-number and the layer-size of each layer is uniquely determined by the pair {*k, P*}. The case of *m*_*last*_ is particularly interesting and will be useful in explaining several of our results. Some of these properties are as follows:

1. For all *P* ≠ 1, *m*_*last*_ is independent of *k* (see Appendix A, Property A.2), *i.e*., it is not affected by the leading zeroes in the binary string of length *k* representing *P*.
2. The value of *m*_*last*_ for any *k*-input NCF with bias *P* = 4*t* + 3 and bias *P* ^*′*^= 4*t* + 5 are equal for any *t* ∈ ℕ_0_ (see Appendix A, Property A.3).
3. All bias *P* (≠1) values of a *k*-input NCF with *m*_*last*_ = *m* can be expressed as *P* = *S*.2^*m*+1^ + 2^*m*^ *±* 1 for some *S* ∈ ℕ_0_ (see Appendix A, Property A.4)

#### 2.3.2. Number of NCFs for a given number of inputs

Li *et al*. [33, 34] provided a formula for calculating the number of NCFs for any number of inputs *k*. Here, we present that formula in our notation. The number of *k*-input NCFs with bias *P* is given by:

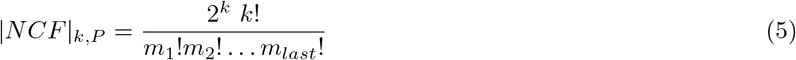

The set {*m*_1_, *m*_2_, …, *m*_*last*_} is uniquely determined for any *k* and *P* as explained in section 2.3.1. The total number of NCFs for a particular *k* can be obtained by summing Eq. (5) over all odd biases as follows:

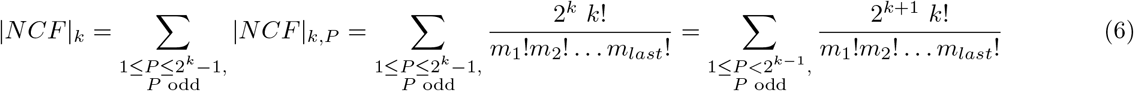

Note that the layer-number and layer-length of each layer is invariant under complementation of the NCF as one can obtain the binary string representation of 2^*k*^ − *P* by replacing the 0 bits by 1 and vice versa in the binary string representation of *P*.

### 2.4. Chain functions

Chain functions were first introduced by Gat-Viks and Shamir [23]. They were found to be a sub-type of NCFs [3, 25]. We provide below 4 equivalent definitions of chain functions: the original definition by Gat-Viks and Shamir [23], the definition of Akutsu *et al*. (which uses the Boolean expression) [25] along with an alternative version, and the definition of Kauffman *et al*. [3] (based on canalyzing inputs).

#### Definition 3 (Gat-Viks and Shamir 2003)

Consider a regulatee *g*_*k*+1_ driven by the ordered inputs **g** = (*g*_1_, …, *g*_*k*_) where each *g*_*i*_ can take on a binary value denoted *x*_*i*_. Introduce a control pattern **c** = (*c*_1_, …, *c*_*k*_) with **c** ∈ {0, 1}^*k*^. Then the associated chain function is defined via the propagation of a so-called *influence* value from one input index to the next as follows. First, the iteration is initialized via *infl*(*g*_1_) = 1. Second, it is continued using the recursion

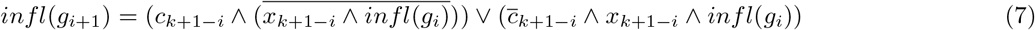

Lastly, the iteration ends by setting the chain function’s output value *x*_*k*+1_ to *infl*(*g*_*k*+1_). Qualitatively, Eq. (7) describes how the control bits are used to go from one input to the next.

#### Definition 4 (Akutsu *et al*. 2011)

A function *f* is a chain function with the control pattern 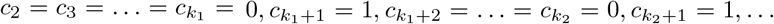 and input variable states of the form *x*_1_, *x*_2_, …, *x*_*k*_ if and only if it has either of the following two forms:

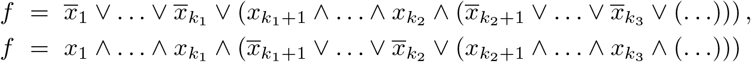

where the former and the latter expressions correspond to *c*_1_ = 1 and *c*_1_ = 0 respectively.

Note that the control bits here produce the same chain function as the ones in the Gat-Viks and Shamir definition and so the rather procedural approach there can be quite simply interpreted by switching to this Akutsu *et al*. framework. This expression points out some unique properties of the chain functions:

1. Every time a control variable takes the value 1, a new layer (for definition of layer see Definition 2) begins.
2. At all but the last layer a positive variable is followed by an AND (∧) operator, whereas a negative variable is followed by an OR (∨) operator.

We now define chain functions using the binary representation of the bias instead of the control pattern in the following manner.

#### Definition 4*.

Consider a bias *P* such that the first *k* − 1 significant bits of the *k*-bit binary representation of *P* are 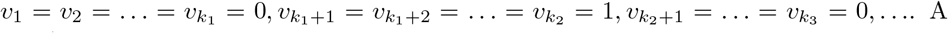 *k*-input BF with the ordered input variables *x*_1_, *x*_2_, …, *x*_*k*_, is a chain function with that bias if and only if:

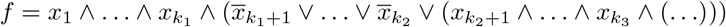

Similarly, when the first *k* − 1 significant bits of the *k*-bit binary representation of *P* are 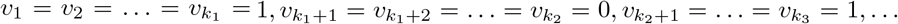, a function will be a chain function with that bias if and only if:

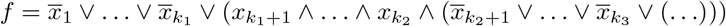

The relationship between the binary representation of the bias *P* and the layer structure of a chain function expression can be understood as follows. Consider a *k*-input chain function with bias *P* (*P* is odd since chain functions are a sub-type of NCFs, which have odd bias). Suppose, the binary representation of *P* is given by:

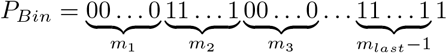

Then, the associated expression of a chain function will be of the following form

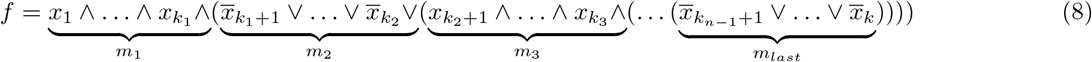

where, *m*_1_ = *k*_1_, *m*_2_ = *k*_2_ − *k*_1_, *m*_3_ = *k*_3_ − *k*_2_, …, and *m*_*last*_ = *k* − *k*_*n*−1_. Note that other similar expressions where exactly one variable in the last layer is negated are also chain functions for the same bias *P*. Let us illustrate this via a specific example of a 6 input chain function with bias 7 with respect to an ordering on its inputs {*x*_1_, *x*_2_, …, *x*_6_}. The 6 bit binary representation of 7 is ‘000111’ and therefore *v*_1_ = *v*_2_ = *v*_3_ = 0 and *v*_4_ = *v*_5_ = 1. Then, the expression of the associated chain function could be any of the following:

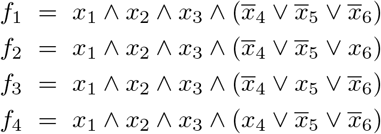

In the last layer, there are *m*_*last*_ variables connected by *m*_*last*_ − 1 operators. Therefore, the ‘last’ variable *x*_6_ which is not followed by any operator can be either positive or negative. Thus, *f*_1_ and *f*_2_ are consistent with the definition of chain functions. Note that *f*_3_ and *f*_4_ in the above equations may appear inconsistent with the definition of chain functions in that the operators associated with the positive variables *x*_5_ and *x*_4_ respectively are both ∨ operators. However, since all the variables in this last layer are connected by the ∨ operator, one is allowed to exchange variables along with their sign within that layer without altering the resulting BF. In this manner, the positive variables may be moved to the last position, thereby restoring the apparent inconsistency between *f*_3_ and *f*_4_.

Lastly, Kauffman *et al*. [3] defined the chain functions as a specific constraint on the canalyzing inputs of the NCFs as follows:

#### Definition 5 (Kauffman *et al*. 2003)

A *k*-input chain function is a *k*-input NCF where the first (*k* − 1) canalyzing input values are 0. The last input is canalyzing in both 0 and 1.

Henceforth, we will refer to these chain functions as *ChF*_0_ since their canalyzing input values are 0. That viewpoint opens up other options for constraining NCFs. In particular, it is natural to consider having the (first *k* − 1) canalyzing input values be 1 instead of 0; we then obtain a new sub-type of NCFs as will now be presented.

#### 2.4.1. *Introduction of Chain-1 Functions (ChF*_1_)

##### Definition 6.

A *k* -input chain-1 function (*ChF*_1_) is a *k* -input NCF where the first *k* − 1 canalyzing input values are 1 and the last input is canalyzing in both 0 and 1.

Just as for *ChF*_0_ functions, one can provide multiple equivalent definitions of chain-1 functions. Here we provide one analogous to that in Definition **4**^*∗*^.

##### Definition 7.

A *k* input chain-1 function with bias *P*, where the first *k* − 1 significant bits of the *k*-b it binary representation of *P* are 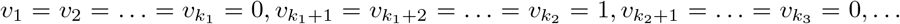, with the ordered input variables *x*_1_, *x*_2_, …, *x*_*k*_ is given by

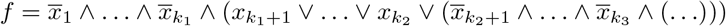

Similarly, when the first *k* − 1 significant bits of the *k*-bit binary representation of *P* are 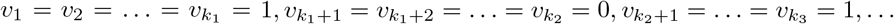, the chain function will be of the form

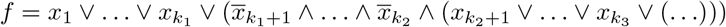

Clearly, the Boolean expressions of *ChF*_1_ are similar to those of *ChF*_0_ and one can obtain a *ChF*_0_ from a *ChF*_1_ and vice versa by either flipping the sign of all the variables (negating all the variables) or flipping all the operators (i.e., replacing ∧ with ∨ and vice versa). By definition, both classes *ChF*_0_ and *ChF*_1_ are subsets of NCF and thus have odd bias. From *k* = 3 onwards, they also form completely disjoint classes (see Appendix B, Property B.1). We will refer to the union of *ChF*_0_ and *ChF*_1_ as **generalized chain functions** or ***ChF***_***U***_.

### 2.5. Description of the reference biological datasets

In this section, we describe the 3 reference biological datasets containing the BFs used to quantify the abundance and enrichment of the *ChF*_0_, *ChF*_1_, *ChF*_*U*_ and *non*–*ChF*_*U*_ NCF types of BFs.

- **BBM benchmark dataset**: This dataset was adapted from the BBM benchmark dataset provided in [26], which consisted of 219 models of Boolean GRNs. Of these, we selected only the manually reconstructed ones, amounting to total 134 models. From those 134 models, we extracted 5990 BFs (regulatory logic rules) when restricting to cases having at most 10 inputs.
- **MCBF dataset**: This dataset was published in [13]. It consists of 2687 BFs recovered from 88 manually reconstructed discrete models of Boolean GRNs. This was downloaded from the Github repository https://github.com/asamallab/MCBF. Here as well we restricted ourselves to a maximum of 10 inputs per BF, leading to 2682 BFs.
- **Harris dataset**: This dataset is published in [3, 24]. It consists of 139 BFs. This dataset has only BFs with 5 or less inputs and has been used exhaustively.

### 2.6. Relative enrichment and associated p-values

Consider a type of BF *T* and one of its sub-types, say *T*_*s*_. In our study we are interested in relative enrichment of the sub-type *T*_*s*_ within its englobing type *T* when considering a particular biological dataset. For instance we can examine the case where *T* = NCF and *T*_*s*_ = *ChF*_1_. We define the relative enrichment for a given number of inputs *k* by *E*_*R*_ = (*f*_*s*,1_*/f*_1_)*/*(*f*_*s*,0_*/f*_0_) where *f*_*s*,1_ and *f*_1_ are the fraction of BFs belonging to *ChF*_1_ and NCF respectively in the considered biological dataset, and *f*_*s*,0_ and *f*_0_ are the fraction of BFs belonging to *ChF*_1_ and NCF respectively when considering all possible such functions. In typical cases the biological dataset will have an over representation of BFs of type *T* and *T*_*s*_ but by considering the relative enrichment we can test the hypothesis that this last enrichment is driven solely by the property of being in *T*; if the relative enrichment of *T*_*s*_ within *T* is statistically inconsistent with the value 1, then we can reject the hypothesis. In particular, if *E*_*R*_ is large, then there must be other factors than “belonging to *T*” driving this relative enrichment. It is possible to compute the *p*-values associated with these relative enrichments to test their statistical significance. The code for computing these *p*-values is available in our GitHub repository.

### 2.7. Reconstructed biological models used to implement model selection

#### 2.7.1. Pancreas cell differentiation model

The Pancreas cell differentiation network [28] is a reconstructed GRN that controls the differentiation of cells in the pancreas. This model consists of 5 genes and 13 edges. This Boolean model has 3 fixed points corresponding to the cell types Exocrine, *β/δ* cell progenitor and *α*/PP cell progenitor (see SI Table S1).

#### 2.7.2. Arabidopsis thaliana root stem cell niche (2010 model)

The RSCN-2010 Boolean model (*model A* in [29]) is a reconstructed Boolean GRN that controls the differentiation of cells in the root stem cell niche (RSCN) of *Arabidopsis thaliana*. The RSCN is located in the root tip of the plant. The 2010 BN has 9 nodes and 19 edges. The BF at each gene of this model is provided in SI Table S2. This Boolean model has 4 fixed points corresponding to the cell types: Quiescent center (QC), Vascular initials (VI), Cortex-Endodermis initials (CEI) and Columella epidermis initials (CEpI) (see SI Table S3). The BF for the AUX node for all our computations derived from this model is the same as the one provided in [35].

#### 2.7.3. Arabidopsis thaliana root stem cell niche (2020 model)

The RSCN-2020 Boolean model [30] is the latest reconstructed Boolean GRN that controls the differentiation of cells in the root stem cell niche (RSCN) of *Arabidopsis thaliana*. The 2020 BN has 18 nodes and 51 edges. The BF at each gene of this model is provided in SI Table S4. This model has 6 biological fixed points corresponding to the cell types: Quiescent center (QC), Cortex-endodermis initials (CEI), Peripheral Pro-vascular initials (P. Pro-vascular PD), Central Pro-vascular initials (C. Pro-vascular PD), Transition domain (C. Pro-vascular TD2) and Columella initials (Columella 1) (see SI Table S5).

### 2.8. Deriving biologically plausible ensembles

We use the methodology developed in [27, 28] to generate a set of biologically plausible ensembles of models. That procedure involved applying several constraints on the truth table: (i) fixed point constraints, (ii) biologically meaningful BFs, (iii) BFs that obey the signs of the interactions in the network architecture. Constraint (i) ensures that all the models in the resulting ensemble recover the expected biological fixed points (fixed points that correspond to the cell states). However, this constraint does not guarantee the absence of spurious attractors. Constraint (ii) restricts the type of BFs to biologically meaningful ones such as NCFs or RoFs [13]. In this work, we impose the *ChF*_0_, *ChF*_1_ and *ChF*_*U*_ functions as our biologically meaningful types of BFs for different biological models. Constraint (iii) forces the imposed biologically meaningful BFs to obey the signs of the interactions that have been observed experimentally.

### 2.9. Relative stability and the mean first passage time

The relative stability of a pair of cell states (attractors) quantifies the propensity to transition from one cell state to the other versus in the opposite direction. Several measures of relative stability have been introduced in the literature [27, 28, 36] of which the Mean First Passage Time (MFPT) captures best the directional aspect of state transitions.

Here we succinctly present the mathematical framework proposed by Zhou *et al*. [28] to define the relative stability of a pair of cell states. To begin, those authors extend the Boolean dynamics to render them stochastic. Mathematically, this change is specified by an *N × N* transition matrix (*N* being the number of nodes in the network):

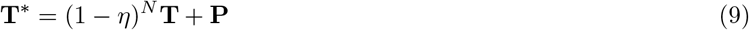

where **T** and **P** are the matrices representing the deterministic and stochastic components of the dynamics respectively. *T*_*lm*_ are the entries of the deterministic matrix **T**, such that *T*_*lm*_ = 1 if updating the state *m* via BFs **F** = {*f*_1_, *f*_2_, …, *f*_*N*_} gives the state *l* (here *l, m* ∈ {0, 1, …, 2^*N*^ − 1} and 0 otherwise). *P*_*lm*_ are the entries of the *perturbation* matrix **P** such that *P*_*lm*_ is the probability that a noise *η* alone drives the transition from state *m* to state *l*. To be explicit, *P*_*lm*_ is defined via:

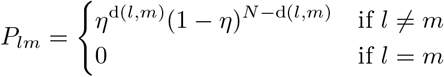

where d(*l, m*) is the Hamming distance between *l* and *m*. In brief, if the noise (*η*) does not alter the state of the network, then one applies the deterministic dynamics, otherwise, noise drives the dynamics.

The number of steps along a state space trajectory starting at state *m* and terminating at the first occurrence of *l* in a stochastic process is called the first passage time from state *m* to *l*. Its average over a large number of trajectories is then the MFPT from *m* to *l* and is denoted by *M*_*lm*_. We use a stochastic method proposed in [27] to compute the MFPT. If *u* and *v* denote 2 biological fixed points (cell states), then the MFPT *M*_*uv*_ is the average of the number of time steps taken over a large number of trajectories starting at state *v* and evolved iteratively under the above-mentioned dynamics till state *u* is reached. Finally, the authors of [28] define the relative stability of cell state *u* compared to cell state *v* via:

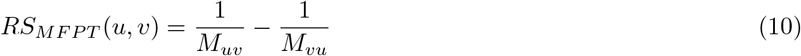

*RS*_*MF P T*_ (*u, v*) *>* 0 if cell state *u* is more stable than cell state *v*. In practice, in this work, the noise intensity parameter value is set to *η* = 0.05; furthermore, 3000 state space trajectories of the first passage times are averaged over to obtain our estimates of each MFPT.

### 2.10. Model selection using relative stability

Having generated ensembles of biologically plausible models by applying the methodology described in section 2.8, it is appropriate to further zero in on a smaller subset of Boolean models by applying more severe biological constraints. Using relative stability constraints derived from the published literature, such a reduction on the ensembles is possible. Provided below are the known relative stability constraints for the 3 different biological models considered here. For the Pancreas cell differentiation model, the 3 relative stability constraints [28] are: (i) Exocrine *< α*/PP progenitor; (ii) Exocrine *< β/δ* cell progenitor; and (iii) *α*/PP progenitor *< β/δ* cell progenitor. For the RSCN-2010 model, the 3 relative stability constraints in [29, 35] are: (i) QC *<* VI; (ii) QC *<* CEI; and (iii) QC *<* CEpI. For the RSCN-2020 model, the 6 relative stability constraints in [27, 30] are: (i) QC *<* CEI/EndodermisPD; (ii) QC *<* P.ProvascularPD; (iii) QC *<* C.ProvascularPD; (iv) QC *<* C.ProvascularTD2; (v) QC *<* Columella1; and (vi) C.ProvascularPD *<* C.ProvascularTD2. The names provided above are the abbreviated names of the cell states, see section 2.7 for their full expansion.

For each of the 3 biological models, we perform an exhaustive search, over their biologically plausible ensembles, for models that obey the above expected relative stability constraints. Additionally, we ‘select’ a model only if the sum over the sizes of the basin of attraction of the biological fixed points for that model is at least as large as that same sum for the original Boolean model to penalize models having spurious attractors.

## 3. RESULTS

### 3.1. Properties of chain-0 and chain-1 functions

We now introduce some properties of the chain-0 (*ChF*_0_) and chain-1 (*ChF*_1_) functions based on the 3 operations on BFs described in section 2.2.3. These properties will be useful in counting the number of *k*-input *ChF*_0_ (or *ChF*_1_) with bias *P* as we shall see in the following section. We explain these properties here for *ChF*_0_ only, but they hold true for *ChF*_1_ as well.

1. A permutation of any chain-0 function is a chain-0 function. Permuting a *ChF*_0_ shuffles the subscripts of the variables in its Boolean expression. Such an operation preserves the sequence of the operators (∧,∨) and of the signs (*x*_*i*_ versus 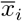) in that Boolean expression, thereby resulting in a *ChF*_0_.
2. Negating variables in a chain-0 function may or may not produce a chain-0 function. A negation operation performed on any set of variables that are not in the last layer of a *ChF*_0_ does not result in a *ChF*_0_ since the signs of the negated variable and the operator following it become inconsistent with the definition of the *ChF*_0_. For the last layer, if variables are linked by ∧ (or ∨) operator, then at least *m*_*last*_ − 1 variables must be positive (or negative). Thus only some negations in that layer may lead to a *ChF*_0_ (see section 2.4).
3. The complement of a chain-0 function is a chain-0 function. The complementation operation simultaneously flips both the signs of all the variables (positive variables become negative and vice versa), and also the conjunction and disjunction operators (∧ operators are replaced by ∨ and vice versa). Such an operation preserves the sign-operator relations that characterize the *ChF*_0_.
4. *ChF*_0_ and *ChF*_1_ form disjoint classes within the NCFs if and only if *k* ≥ 3 (see Appendix B, Property B.1 for proof, see Fig. 2(a) for illustration).

**FIG. 1.**
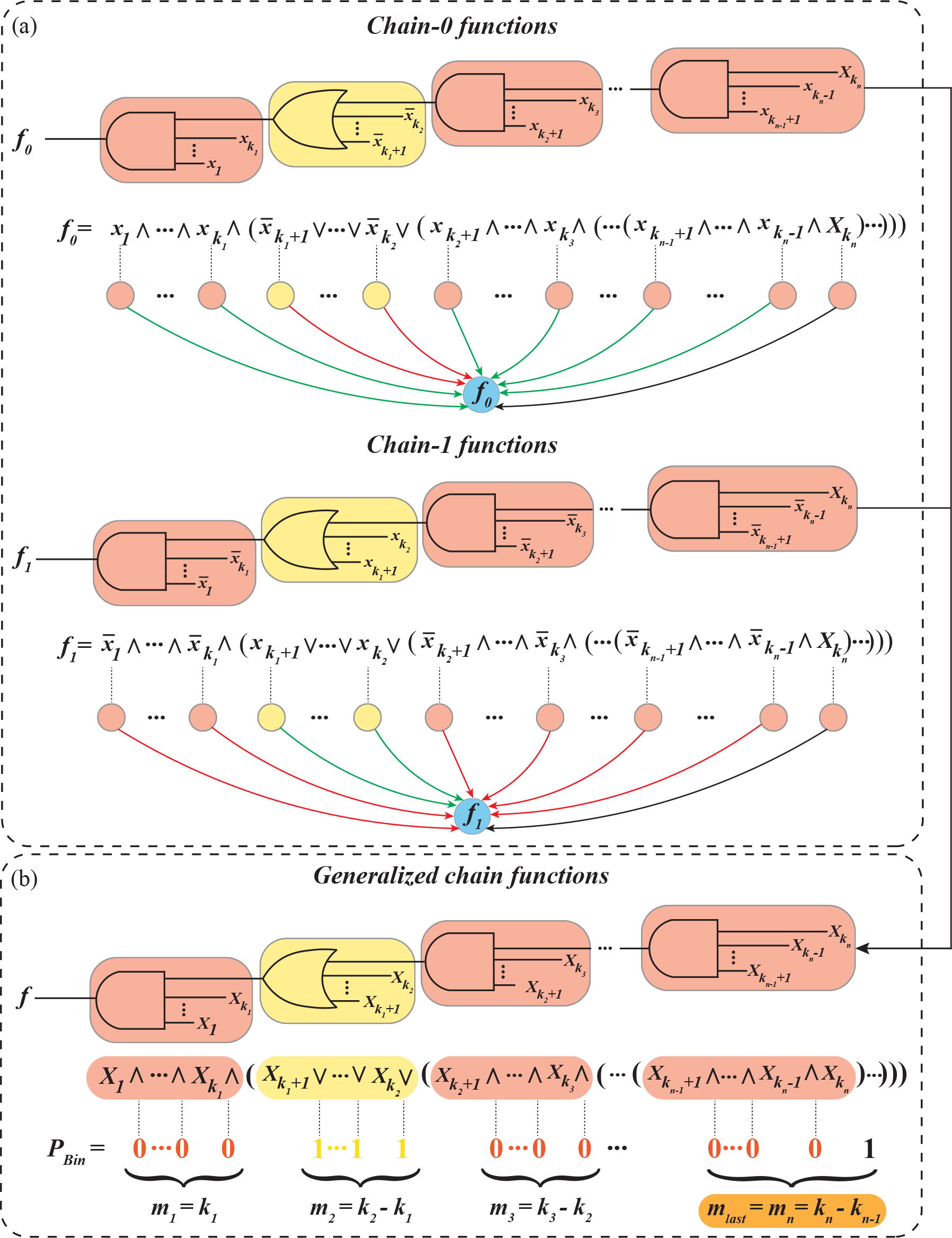
Relationship between layers, operators, variable signs and Boolean string representation of bias. The two circuit diagrams in **(a)** correspond to chain-0 (*f*_0_) and chain-1 (*f*_1_) functions respectively with bias *P <* 2^*k*−1^ and with a odd number of layers, where *k* = *k*_*n*_. Rounded rectangular boxes with alternating colors correspond to successive layers. The schematics below the expressions of *f*_0_ and *f*_1_ denote the interactions driving the output of the BF. Here, the ‘green’ and ‘red’ arrows indicate ‘activatory’ (positive) and ‘inhibitory’ (negative) regulation respectively. The black arrows indicate regulation of either nature. Top part of **(b)** corresponds to the logic circuit diagram of the generalized chain function with bias *P <* 2^*k*−1^ and with an odd number of layers. Here 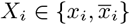 for all *i* ∈ {1, 2, …, *k*_*n*_} with the constraint that the sign of all the variables in a given colored layer be the same (but altering from one layer to the next), with the exception of last layer. The Boolean expression form of the function is provided below the circuit diagram and layers are colored accordingly. The associations of operators and binary representation of the bias (*P*_*Bin*_) are shown (∧ with 0 and ∨ with 1). *m*_*i*_ correspond to the layer-size of the *i*^*th*^ layer for all *i* ∈ {1, 2, …, *k*_*n*_}.

**FIG. 2.**
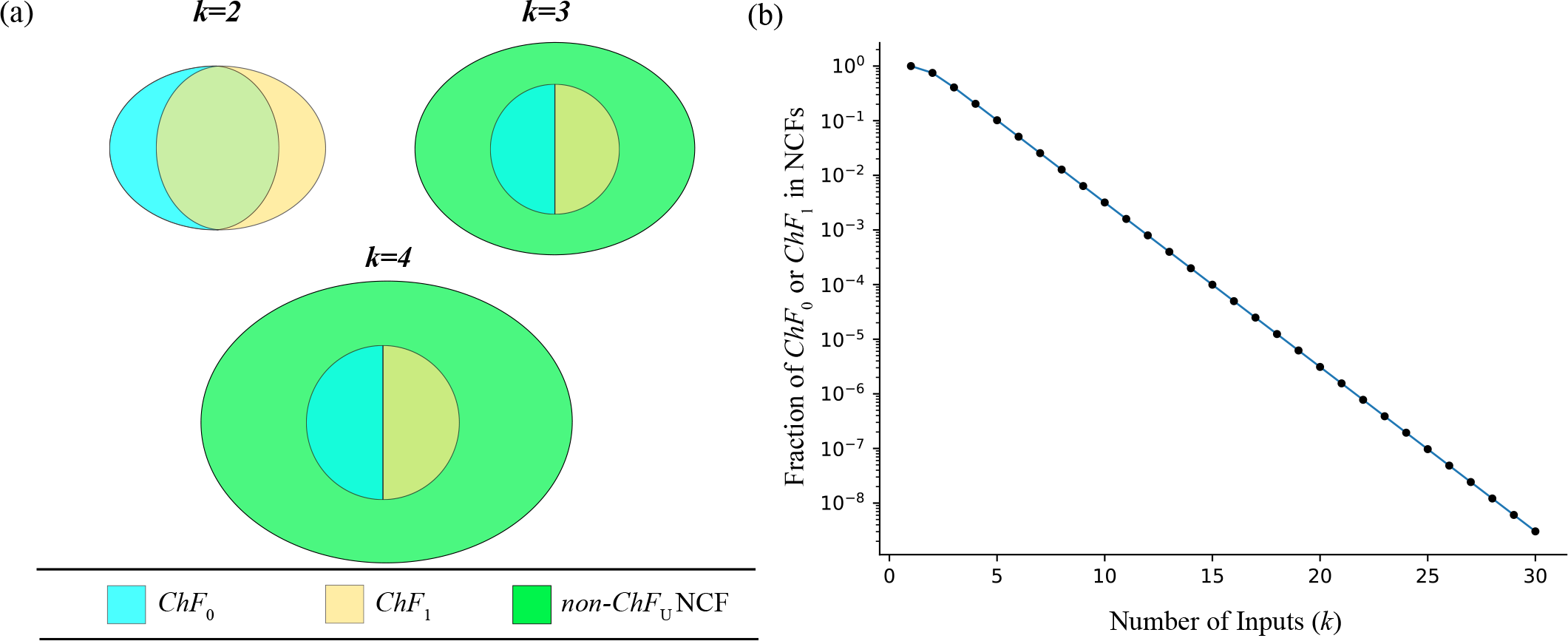
Fraction of chain-0 (or chain-1) functions within NCF for different number of inputs. **(a)** Venn diagram of the space of *ChF*_0_, *ChF*_1_ within the NCFs. *k* = 1 is not shown as *ChF*_0_, *ChF*_1_, *ChF*_*U*_ and NCFs are identical. For *k* = 2, the *ChF*_0_ and *ChF*_1_ do not completely overlap, and all NCFs are *ChF*_*U*_s. For *k* ≥ 3 the *ChF*_0_ and *ChF*_1_ sets become disjoint. **(b)** The fraction of *ChF*_0_ (or *ChF*_1_) within NCF (*y*-axis) is plotted as a function of the number of inputs *k* (*x*-axis). The *y*-axis has a logarithmic scale. The linear trend suggests that the fraction of *ChF*_0_ (or *ChF*_1_) in NCF is an exponentially decaying function of *k*.

### 3.2. Counting the number of chain-0, chain-1 and generalized chain functions

Gat-Viks and Shamir have provided a formula to count the number of *ChF*_0_ for a given number of inputs based on a recursive approach [23]. We take a different approach and first count the number of *k*-input *ChF*_0_ at bias *P*, using which we count the number of *ChF*_0_ by summing over all odd biases. To count the number of *k*-input *ChF*_0_ for a given bias *P*, we need to compute all permutations and negations of a reference chain-0 function that yield all the distinct *ChF*_0_ with bias *P* (this suffices to obtain all *ChF*_0_ with bias *P* as all NCFs with bias *P* form a single equivalence class under permutations and negations of variables [13]). The number of ways to permute *k* variables in Eq. (8) that lead to distinct *ChF*_0_ is the multinomial coefficient 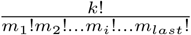, where *m*_*i*_ is the layer-size of the *i*^*th*^ layer. The number of negations (including the identity operation) of a *ChF*_0_ that lead to distinct *ChF*_0_s is 1 + *m*_*last*_. Now for each permutation there are 1 + *m*_*last*_ possible negations, therefore the number of *k*-input *ChF*_0_ for a given bias *P* is:

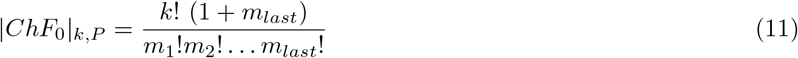

Using Eq. (11), the total number of *k*-input *ChF*_0_ is given by,

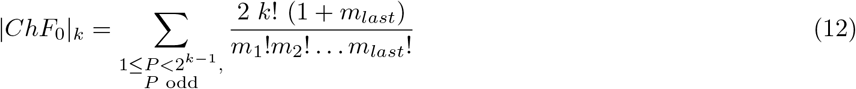

*P* is odd because *ChF*_0_ have odd bias. Furthermore, the factor 2 in this equation accounts for a complementary function with bias 2^*k*^ − *P* associated with each *ChF*_0_ with bias *P <* 2^*k*−1^ since the layer-number and layer-size are invariant under complementation for any NCF. The total number of BFs belonging to the class *ChF*_*U*_ for *k* ≥ 3 is then given by:

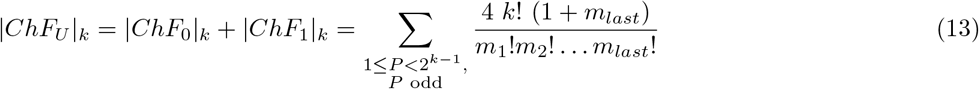

since |*ChF*_0_|_*k*_ = |*ChF*_1_|_*k*_ (see Appendix B, Property B.2) and for *k* ≥ 3, *ChF*_0_ and *ChF*_1_ form disjoint sets. Note, for *k* ≤ 2 (see Appendix B, Property B.1),

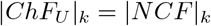

### 3.3. The fraction of chain-0 and chain-1 within NCFs decreases exponentially with the number of inputs

The fraction of NCFs occupied by *ChF*_0_ for different number of inputs is yet to be explored systematically. To begin, we compute the fraction of NCFs that are in *ChF*_0_ for *k* ∈ {1, 2, …, 30} using exhaustive enumeration. Strikingly, this fraction appears to diminish exponentially as *k* grows as shown by the linear trend in the semi-log plot in Fig. 2(b). Furthermore, the generalized chain functions (*ChF*_*U*_) also show this trend since the cardinality of that class of functions is a factor 2 larger than that of *ChF*_0_ for all *k* ≥ 3 (see Appendix B, Property B.2). The fraction of NCFs belonging to a *k*-input *ChF*_0_ can be written in the following form (see Appendix C for derivation):

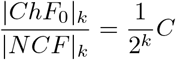

where

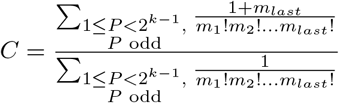

Since *m*_*last*_ ≥ 2 for all *k* ≥ 2, one has *C* ≥ 3. Since *m*_*last*_ is at most *k* for any *k, C* ≤ *k* + 1. From this it follows that the fraction 1*/*2^*k*^ accounts for the observed exponential decay. Exhaustive computation up to *k* = 30 suggests that *C* converges at large *k* towards a value ≈ 3.2588913532709. As the cardinality of *ChF*_0_ is equal to that of *ChF*_1_ (see Appendix B, Property B.2), all the results obtained for *ChF*_0_ are equally valid for *ChF*_1_. Note that this exponential decay has important implications for restricting the space of biologically plausible models as we will demonstrate in later sections.

### 3.4. Bias-wise fractions of chain-0 and chain-1 functions within NCF

To study the behaviour of how the fractions of *ChF*_0_ (or *ChF*_1_) and NCFs within all BFs, and with respect to one another, vary with bias (*P*) for a given number of inputs (*k*), we plot the fraction of: (i) NCFs within all BFs (see SI Fig. S1); (ii) *ChF*_0_ within all BFs (see SI Fig. S2); and (iii) *ChF*_0_ within NCFs (see Fig. 3). Of these, the fraction *ChF*_0_ within NCFs (see Fig. 3) showed several interesting features. Note that since the cardinality of *ChF*_0_ is equal to that of *ChF*_1_ (see Appendix B, Property B.2) for a given *k* and *P*, all results obtained for *ChF*_0_ are equally valid for *ChF*_1_. We list below our observations and will explain them with the following simple and elegant formula derived using Eqs. (5) and (11).

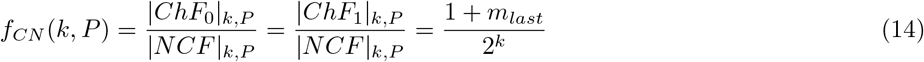

**FIG. 3.**
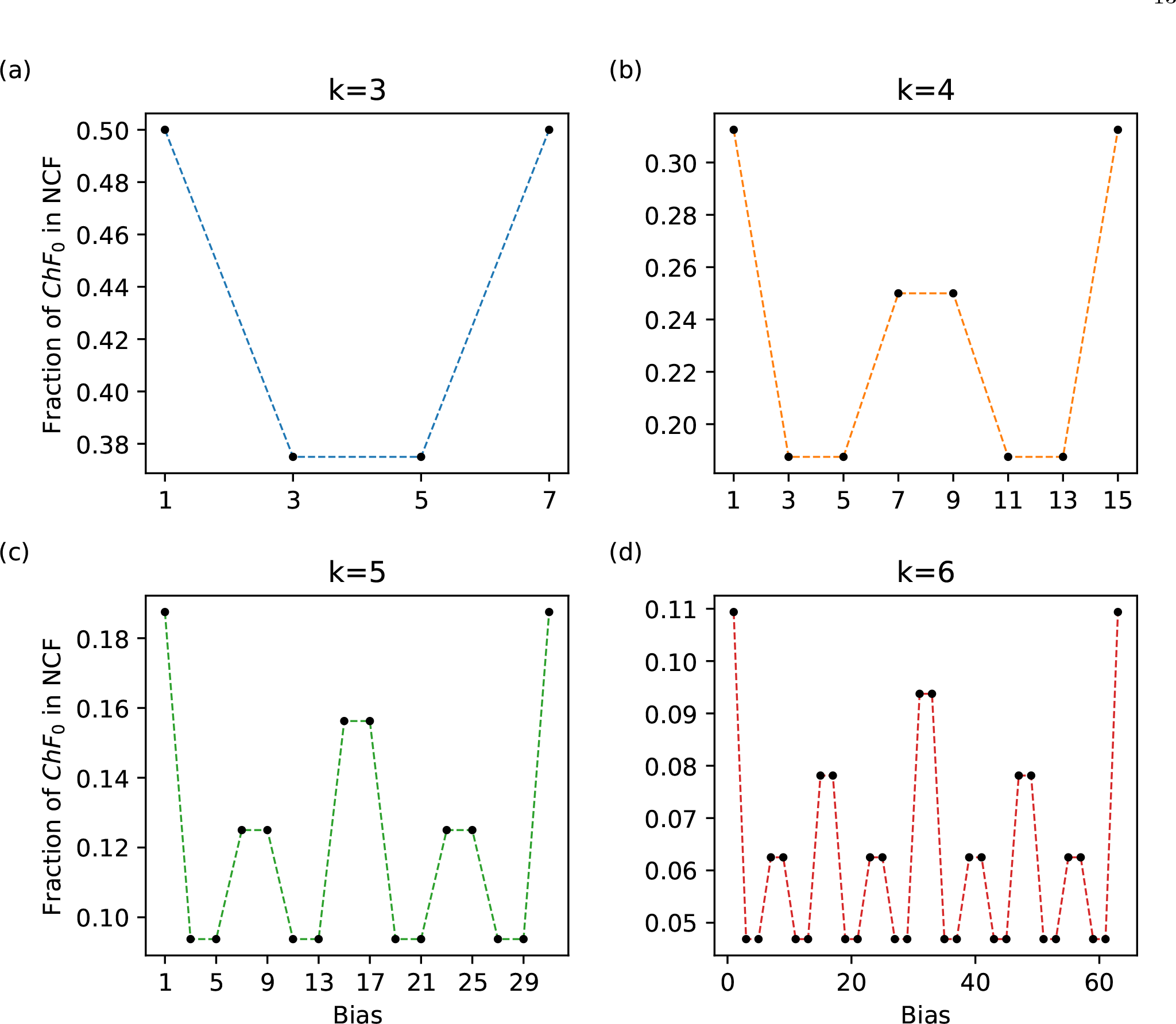
Bias-wise fraction of *ChF*_0_ (or *ChF*_1_) within NCFs. For a given number of inputs (*k*), the fraction of *ChF*_0_ (or *ChF*_1_) in NCFs (*y*-axis) is plotted as a function of the bias (1 ≤ *P* ≤ 2^*k*^ − 1 for odd *P*) (*x*-axis). Subplots correspond to different number of inputs *k* = 3, 4, 5 and 6.

The observations are as follows:

1. *f*_*CN*_ (*k, P*) = *f*_*CN*_ (*k, P* + 2) whenever *P* = 4*t* + 3 for any *t* ∈ ℕ_0_. In other words, those particular pairs of consecutive biases (starting from *P* = 3) have equal *f*_*CN*_ (*k, P*). This implies that for any *k*, the *m*_*last*_ for consecutive biases (see Eq. (14)) are equal. This is indeed the case as we prove in Appendix A, Property A.3.
2. Since *m*_*last*_ determines *f*_*CN*_ (*k, P*), biases with equal *m*_*last*_ also have equal *f*_*CN*_ (*k, P*). For example, using Property A.4 in Appendix A, *m*_*last*_ = 2 for *P* = 3, 5, 11, 13, 19, 21…. This explains why several biases in Fig. 3 have the same *f*_*CN*_ (*k, P*).
3. *f*_*CN*_ (*k, P*) is maximum when *P* = 1 or 2^*k*^ − 1 for a given *k* and it is minimum when *P* = 3, 5, 11, 13, 19, 21, …. For *k* ≥ 2, we know that 2 ≤ *m*_*last*_ ≤ *k*. So, *f*_*CN*_ (*k, P*) is maximum (respectively minimum) when *m*_*last*_ = *k* (respectively *m*_*last*_ = 2). *m*_*last*_ = *k* iff *P* = 1 or *P* = 2^*k*^ − 1 since *m*_*last*_ = *k* corresponds to a *ChF*_0_ (or *ChF*_1_) with a single layer. The maximum value of *f*_*CN*_ (*k, P*) is then (1 + *k*)*/*2^*k*^. However *m*_*last*_ = 2 for several values of *P* (see Appendix A, Property A.4). The minimum value of *f*_*CN*_ (*k, P*) is (1 + 2)*/*2^*k*^ = 3*/*2^*k*^ (when *k* ≥ 2) and (1 + 1)*/*2^1^ = 1 (when *k* = 1).
4. 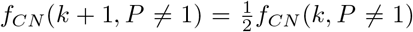 and 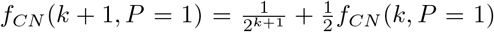. From Eq. (14), it is easy to see that 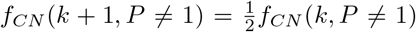 since *m*_*last*_ for *P* ≠ 1 remains invariant for any *k* (see Appendix A, Property A.2). However, when *P* = 1, *m*_*last*_ = *k* and so *f*_*CN*_ (*k, P* = 1) = (1 + *k*)*/*2^*k*^. Replacing *k* by *k* + 1 in this equation, 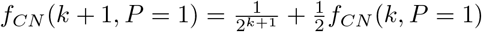.

### 3.5. Preponderance and enrichment of chain-0, chain-1 and generalized chain functions in various reference biological datasets

#### 3.5.1. BBM benchmark dataset

We now shift our focus to quantifying the preponderance of *ChF*_0_, *ChF*_1_ and *ChF*_*U*_ in various biological datasets of regulatory logic rules. We first consider the case of the BBM benchmark dataset [26]. When we quantify the fraction of odd and even bias BFs for a given number of inputs (*k*), we find that the fraction of BFs with odd bias are overwhelmingly larger compared to the fraction of BFs with even bias (see Fig. 4(a)). Next, we compute the fraction of NCFs for each *k* and find that it is enriched for all *k* as shown in Fig. 4(b). Both of these observations are in line with previous studies on the fraction of odd bias BFs and the fraction of NCFs in reference biological datasets [13]. Lastly, for a *k*-input BF belonging to the sub-types of NCFs, namely, *ChF*_0_, *ChF*_1_, *ChF*_*U*_ and *non*–*ChF*_*U*_ NCF, we compute their fraction within the NCFs (shown by the bars in Fig. 4(c) and SI Table S6), their relative enrichments within the NCFs (see SI Table S7) and the associated statistical significance of those enrichments (see *∗* in Fig. 4(c) and SI Table S8 for exact values). We find that *ChF*_1_ is relatively enriched for all values of *k* except at *k* = 2. However, this is not the case for *ChF*_0_ which is not enriched for several values of *k* (*k* = 3, 4, 5 and 9). The union of these two types, *ChF*_*U*_, is significantly enriched for all *k* ≥ 2. These results suggest that the generalized chain function, *ChF*_*U*_, consisting of both *ChF*_0_ and *ChF*_1_ types perhaps constitutes a biologically more meaningful type than either the *ChF*_0_ or *ChF*_1_ separately. Note that at *k* = 1 since all NCFs are also generalized chain functions, it is meaningless to compute a relative enrichment or statistical significance.

**FIG. 4.**
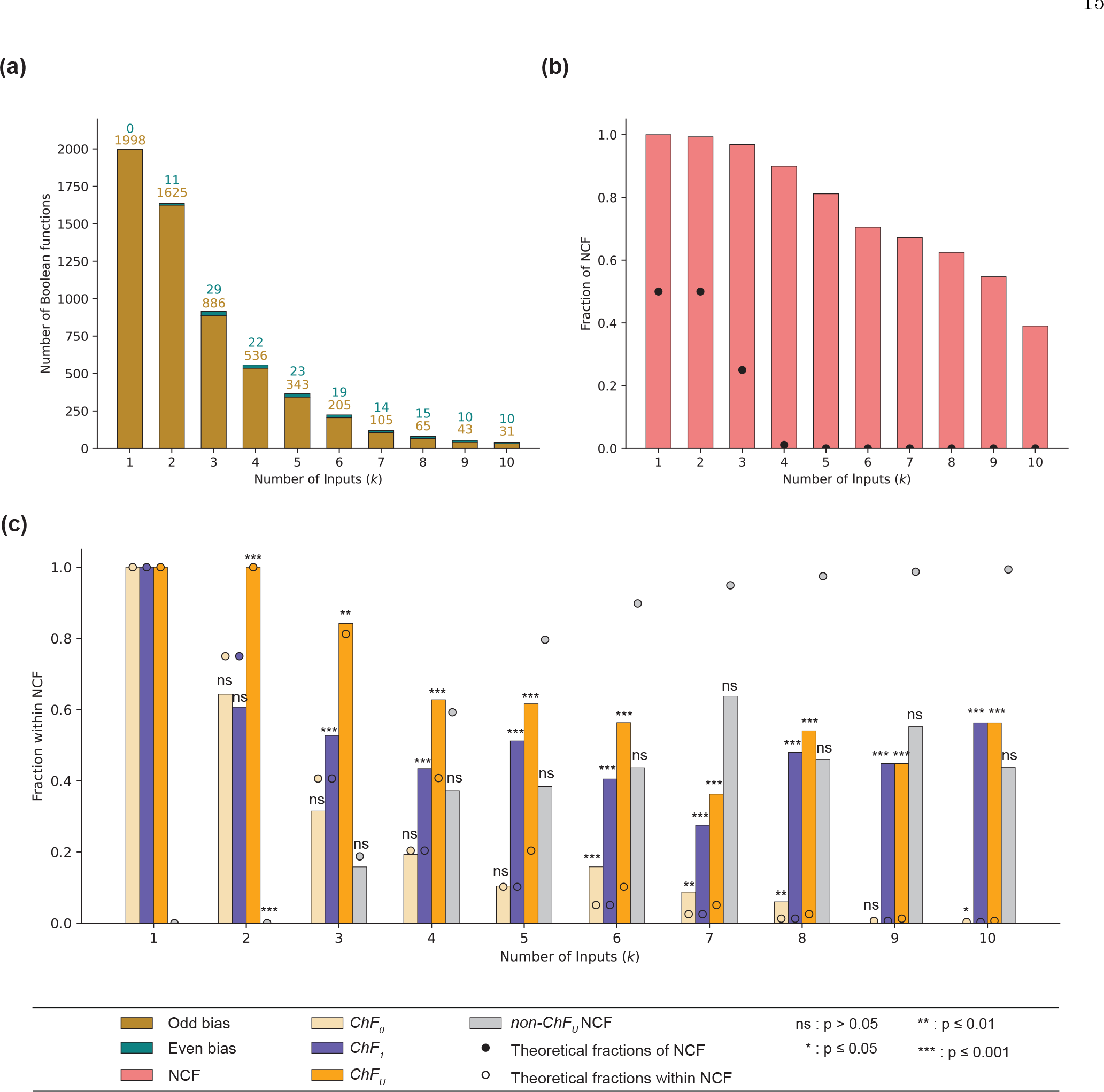
Fractions of *ChF*_0_, *ChF*_1_, *ChF*_*U*_ and *non* − *ChF*_*U*_ NCF within NCFs in the BBM benchmark dataset. **(a)** In-degree distribution of BFs in the BBM benchmark dataset up to *k* ≤ 10 inputs. The frequencies of odd bias are much larger than the frequencies of even bias. **(b)** The fraction of NCFs in the dataset for various number of inputs. The enrichment of NCFs (within all BFs) for any number of inputs is very large and also statistically significant in the BBM benchmark dataset. **(c)** This sub-figure shows the fractions of *ChF*_0_, *ChF*_1_, *ChF*_*U*_ and *non* − *ChF*_*U*_ NCF within NCFs, in theory and in the BBM benchmark dataset as dots and colored bars respectively. The relative enrichments of *ChF*_1_s and *ChF*_*U*_s within NCFs are statistically significant for *k >* 2.

#### 3.5.2. MCBF and Harris datasets

Repeating the previous set of analyses for the MCBF dataset, we find that for a given number of inputs, the fraction of the *ChF*_1_ type within NCFs is typically larger than that of the *ChF*_0_ and *non*–*ChF*_*U*_ NCF types within the NCFs (see SI Fig. S3(a) and SI Table S9). In fact, *ChF*_1_ is relatively enriched within NCFs for all values of *k* except *k* = 2, whereas *ChF*_0_ is relatively enriched only when *k* = 6, 7, 8 (see SI Fig. S3(a) and SI Table S10). Furthermore, *ChF*_*U*_ is relatively enriched within NCFs for all *k* ≥ 3, but only those enrichments where *k >* 3 are statistically significant. The complete set of values for the statistical significance associated with the relative enrichments is provided in SI Table S11. These results obtained using the MCBF dataset are largely in agreement with those of the BBM benchmark dataset.

We now revisit an older dataset of BFs, namely the one published by Harris *et al*. [3, 24]. In this dataset, we find that for any given number of inputs, the fraction of *ChF*_0_ within NCFs is larger than the corresponding fractions of *ChF*_1_ and *non*–*ChF*_*U*_ NCF types (see SI Fig. S3(b) and SI Table S12). Furthermore, the *ChF*_0_ BFs are relatively enriched within NCFs and the enrichments are statistically significant (see SI Tables S13 and S14 respectively). *ChF*_1_ on the other hand is not relatively enriched within NCFs. The generalized chain functions, *ChF*_*U*_, are enriched for *k* = 3, 4, 5 but those enrichments are statistically significant only for *k* = 4, 5. These results are not so concordant with those arising from the BBM benchmark dataset in which the relative enrichment of *ChF*_0_ is not statistically significant for *k* = 3, 4, 5, whereas it is so for *ChF*_1_. It is appropriate to note here that Kauffman *et al*. [3] had observed that a significant fraction of the BFs in that dataset were NCFs and that most of those NCFs were *ChF*_0_. In sum, the generalized chain functions (or *ChF*_*U*_) comprise a special sub-type of NCFs that are quite consistently relatively enriched within NCFs across distinct reference biological datasets and are thereby able to reconcile the discrepancies observed in the relative enrichments of *ChF*_0_ and *ChF*_1_ within NCFs therein.

### 3.6. Model selection using generalized chain functions

In this section, we explore the implications of using generalized chain functions as a constraint in a model selection framework. To do so, we consider 3 biological models, one Pancreas cell differentiation model and two *Arabidopsis thaliana* root development models (see section 2.7 for more details).

First, we apply the framework described in section 2.8 using the generalized chain functions (*ChF*_*U*_) as our biologically meaningful type of BF. The number of allowed BFs at each node of these models (pancreas cell differentiation, RSCN-2010, RSCN-2020) are given in SI Table 15, Table 1 and SI Table 16 respectively. The numbers of models obtained for these three cases are 3600, 60 and 645120 respectively. We remark here that in the RSCN-2020 GRN, no *ChF*_*U*_ satisfied both fixed point and sign conforming constraints at the ARF10 node and so for that node we restricted our choice to NCFs, leading to 4 BFs at ARF10. Note that if we restrict our choice of BFs to NCFs for all nodes in all 3 biological models, we would have got 3600, 1275 and 25019245440 (2.5 *×* 10^10^). Clearly, the resulting number of biologically plausible models is severely reduced (particularly for models having nodes with large number of inputs) compared to the case where NCFs are used as a constraint. This allows us to *exhaustively* search these ensembles for models that satisfy the relative stability constraints provided in section 2.10.

**TABLE 1.** Nodewise enumeration of the number of BFs that satisfy biological constraints for the RSCN-2010 GRN. ‘Nodes’ are the names of the nodes in the network and *k* is the associated number of inputs to that node. The columns *ChF*_0_, *ChF*_1_ and *ChF*_*U*_ give the number of chain-0, chain-1 and generalized chain functions that satisfy biological fixed point constraints and sign conforming constraints at each node of the RSCN-2010 GRN. Imposing *ChF*_*U*_ leads to 60 Boolean models.

| Nodes | $k$ | $ChF_0$ | $ChF_1$ | $ChF_U$ |
| --- | --- | --- | --- | --- |
| PLT | 1 | 1 | 1 | 1 |
| AUXIN | 1 | 1 | 1 | 1 |
| ARF | 1 | 1 | 1 | 1 |
| AUXIAA | 1 | 1 | 1 | 1 |
| SHR | 1 | 1 | 1 | 1 |
| SCR | 4 | 5 | 0 | 5 |
| JKD | 2 | 1 | 0 | 1 |
| MGP | 3 | 1 | 0 | 1 |
| WOX5 | 5 | 12 | 0 | 12 |

Next, we impose known relative stability constraints to shrink the space of biologically plausible models further. Using the MFPT as our relative stability measure, we find that: (i) 19 models satisfy the expected hierarchies for the Pancreas cell differentiation GRN with no spurious attractors. (ii) 16 models satisfy the expected hierarchies for the RSCN-2010 GRN with no spurious attractors. (iii) For the RSCN-2020 GRN, no model satisfied all the expected hierarchies. However, we found that 82355 models violated exactly one hierarchy, namely, ‘C.ProvascularPD *<* C.ProvascularTD2’ of which 28453 did not have any spurious attractors. This result is a major improvement over the original Boolean model proposed by [30] which violated 2 expected hierarchies (‘C.ProvascularPD *<* C.ProvascularTD2’ and ‘QC *<* CEI/EndodermisPD’) and furthermore had a spurious cyclic attractor. In all, for the RSCN-2020 GRN, about 4.4% of models that formed our biologically plausible ensemble using *ChF*_*U*_ yielded improved models.

We have thus demonstrated the utility of generalized chain functions as a biologically meaningful type that can severely restrict the space of biologically plausible models and yet can yield models that satisfy conditions on relative stability.

## 4. DISCUSSION AND CONCLUSIONS

In this work, we have addressed the question of whether there are certain sub-types of NCFs that drive the enrichment of the NCFs in biological regulatory network models. Starting with a known sub-type of NCF, namely the chain functions (or chain-0 functions), we propose two other types, specifically, its dual class – the chain-1 functions, and its union with the chain-1 functions, the generalized chain functions. We first derive an analytical formula to count these functions for a given number of inputs and a given bias. Using this we show that the fraction of chain-0 (or chain-1) functions decreases exponentially within NCFs as the number of inputs increases. Furthermore, our formula can explain several features of the fractal-like pattern observed between the fraction of chain-0 (or chain-1) functions in NCFs and the bias, for a fixed number of inputs. We then test for enrichment of the chain-0, chain-1 and generalized chain function within NCFs in a large dataset of reconstructed Boolean models (the BBM benchmark dataset): the result is that generalized chain functions are indeed highly enriched. In fact, using 2 other datasets of regulatory logics, namely, the MCBF and the Harris dataset, the same result holds. In addition, we demonstrate how generalized chain functions can severely constrain the space of biologically plausible models using 3 different biological models. Lastly, we perform model selection on those models using known relative stability constraints and are able to zero-in on a smaller subset of models that are more biologically plausible.

Since their introduction by Gat-Viks and Shamir [23], the chain-0 functions have hardly received any attention. Kauffman *et al*. identified that chain-0 functions were the sub-type of NCFs for which all the canalyzing input values are 0 and those authors showed its preponderance in the dataset of Harris *et al*. [24]. Akutsu *et al*. [25] put forward the Boolean expression framework associated with chain-0 functions [25], making explicit the dependence of logical operators (∧, ∨) on the signs of the variables preceding it. Surprisingly, its dual class had never been proposed nor explored as a potentially biologically relevant type. Our work builds on all these works, piecing together several concepts pertaining to various representation of chain functions in a systematic manner, thereby providing insights into the nature of chain-0 (and chain-1) functions and the space they occupy within NCFs. Why are generalized chain functions preponderant in biological datasets? For one, the justifications for an enrichment of chain-0 functions given by Gat-Viks and Shamir [23] also apply to chain-1 functions. Furthermore, we may speculate that a somewhat qualitative justification lies in that generalized chain functions have lower complexity than general NCFs when using the framework of Kauffman *et al*., because of the very simple dependence of the operator following a variable and its sign (except for the last variable). The caveat here is that having that simpler description is dependent on the way one represents the functions, so for instance when using the representations of Gat-Vits and Shamir or of Akutsu *et al*., the aforementioned simplicity is no longer apparent. One may also speculate that rather than being subject to a direct selection (e.g., for simplicity,) chain functions are selected for indirectly, likely through their possible modulation of network dynamics. Indeed, network dynamics are subject to strong selection pressures, be-it for robustness or evolvability.

Although these results provide novel insights, it is appropriate to make explicit some limitations in this work. Firstly, we have seen that chain-0 functions were enriched in the Harris dataset, however, when considering much larger datasets such as the BBM benchmark dataset and MCBF dataset, there are multiple values of the number of inputs where there is no such enrichment. Similarly, the chain-1 functions are enriched in the BBM benchmark dataset and MCBF dataset, whereas they are not enriched in the Harris dataset. With more data, we may expect that several other sub-types of NCFs may also contribute to the enrichment of NCFs. Furthermore, there is likely to be an (un)conscious subjective bias on BF logic when reconstructing Boolean models, and so we cannot exclude that such a bias is responsible for the enrichments found. Secondly, the generalized chain functions can be so severely restrictive that it may be impossible to find corresponding BFs that satisfy both the fixed point and sign-conforming constraints as we found in the RSCN-2020 model.

Our findings also invite several directions for future work. One is to quantify properties of the *generalized chain functions* that distinguish them from other NCFs and are likely responsible for their being so prevalent in these biological reference datasets. Another is to analytically tackle the question of whether the ratio of the number of the chain functions to that of the NCFs for a large number of inputs follows a rigorous exponential behaviour.

## AUTHOR CONTRIBUTIONS

Designed the research: S.M., P.S., Aj.S., O.C.M., Ar.S.; Performed the research: S.M., P.S., Aj.S., O.C.M., Ar.S.; Performed the computations: S.M., P.S., Aj.S.; Wrote the paper: S.M., P.S., Aj.S., O.C.M., Ar.S.

## ACKNOWLEDGMENTS

Areejit Samal acknowledges support from the Max Planck Society, Germany, through the award of a Max Planck Partner Group in Mathematical Biology, and the Department of Atomic Energy (DAE), Government of India, through the Apex project to The Institute of Mathematical Sciences (IMSc), Chennai. IPS2 benefits from the support of Saclay Plant Sciences-SPS (ANR-17-EUR-0007).

## DATA AND CODE AVAILABILITY

All data and codes needed to reproduce the results in this manuscript are deposited in GitHub and are available at: https://github.com/asamallab/GenChF.

## Appendix A: Relationship between bias, logical operators and layers in NCFs

### Property A.1.

For any *k*-input NCF, the bias *P* determines the operator sequence in its Boolean expression.

*Proof*. It is evident from the definition of NCF (see Definition 1) that the value of *b*_*i*_ determines the operator following *x*_*σ*(*i*)_ for *i* ≤ *k* − 1. Put simply, if *b*_*i*_ is 0 or 1, the operator following *x*_*σ*(*i*)_ should be ∧ or ∨ respectively. So, there is a clear relation between the sequence of *b*_*i*_ and the sequence of operators. Now, the value *b*_*i*_ fills up 2^*k*−*i*^ rows of the truth table of the NCF. Hence, the bias (*P*) of a NCF can be written in the following form,

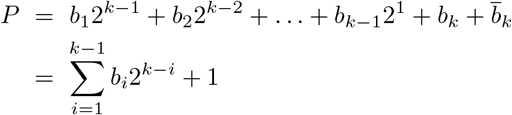

Thus the binary representation of the bias *P* is given by *P*_*Bin*_ = *b*_1_*b*_2_ … *b*_*k*−1_1. This proves that from the binary representation of *P* we can obtain the operator sequence of a NCF with bias *P* such that the bits 0 and 1 encode the operators ∧ and ∨ respectively while considering the sequence from left to right. □

### Property A.2.

For all odd *P* (*P* ≠ 1), *m*_*last*_ is independent of *k* for any *k*-input NCF.

*Proof*. We can obtain the (*k* + 1)-bit binary representation of a (*k* + 1)-input NCF with bias *P* by adding a 0 to the most significant end of the *k* -bit binary representation of bias *P*. More succintly, *P*_*Bin*_(*k* + 1) = ‘0*P*_*Bin*_(*k*)’. This does not alter the value of *m*_*last*_ for all possible bias values except *P* = 1. When *P* = 1, adding a 0 at the most significant end increases the *m*_*last*_ by 1 since *P* = 1 corresponds to a single layer NCF where all but the least significant bit is 0. □

### Property A.3.

The values of *m*_*last*_ for *k*-input NCFs with bias *P* = 4*t* + 3 and bias *P* ^*′*^= 4*t* + 5 are equal, for any *t* ∈ ℕ_0_.

*Proof*. We want to prove that, the values of the *m*_*last*_ for the biases 3, 7, 11, 15, 19, … will be equal to the values of the *m*_*last*_ for the biases 5, 9, 13, 17, 21, … respectively for any *k*-input NCF. Since the biases of the first category are of the form 4*t* + 3 for *t* ∈ ℕ_0_, it becomes evident that the binary representations of these biases will have 1 as the bit to the left of the least significant bit. Consequently, it becomes apparent that their last layer is composed of 1 s (or equivalently of ‘∨’ operators). Therefore, their general binary form can be expressed as:

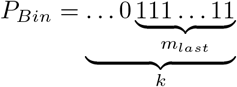

Now, the next odd bias (each of which is of the form 4*t* + 5 for *t* ∈ ℕ_0_) is obtained by adding 2 to *P*, which is the same as performing the Boolean addition *P*_*Bin*_ + 00 … 0010. This yields the following result:

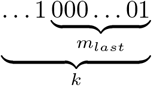

Hence, the value of *m*_*last*_ will remain the same for a bias *P* = 4*t* + 3 and *P* ^*′*^ = 4*t* + 5, for any given value of *t* ∈ ℕ_0_ □

### Property A.4.

All bias *P (*≠ 1) of a *k*-input NCF with *m*_*last*_ = *m* can be expressed as *P* = *S*.2^*m*+1^ + 2^*m*^ *±* 1 for some *S* ∈ ℕ_0_.

*Proof*. It is easy to see that the smallest possible value of *P* for which we get *m*_*last*_ = *m* for any *m* will have the following binary representation

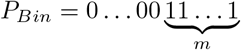

So, the integer form of the bias *P* is given by

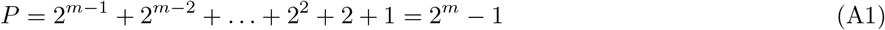

Again *P* can be expressed as

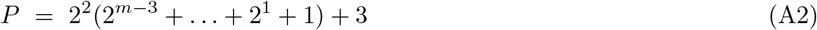

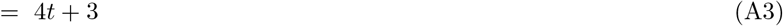

where, *t* = 2^*m*−3^ + … + 2^1^ + 1. So, from Property A.3, for the bias *P* + 2 = 2^*m*^ + 1 also, the value of *m*_*last*_ = *m*. So, given *m*_*last*_ = *m*, the associated two lowest values of *P* have the binary form

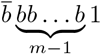

where *b* ∈ {0, 1} which takes the form 2^*m*^ + 1 and 2^*m*^ − 1 (when converted to the integer form) upon taking *b* as 0 and 1 respectively. Now given the binary form 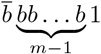 one is free to add any number of bits to the left and *m*_*last*_ will still remain equal to *m*. In other words, any bias *P* ^*′*^ with the binary representation 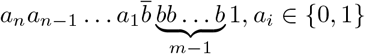 will have *m*_*last*_ = *m, ∀n* ∈ ℕ. We now express *P′* in its integer form as follows:

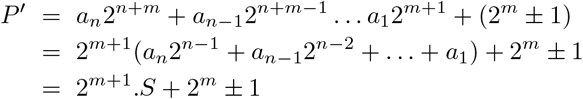

where *S* ∈ ℕ can be expressed in the form *a*_*n*_2^*n*−1^ + *a*_*n*−1_2^*n*−2^ + … + *a*_1_ with suitable choices of *a*_*i*_’s and *n*. □

## Appendix B: Properties of *ChF*_0_ and *ChF*_1_

### Property B.1.

For *k* ≥ 3, the two sub-types of NCF namely *ChF*_0_ and *ChF*_1_ forms disjoint classes.

*Proof*. Let us discuss the scenario for different values of *k*.

- ***k* = 1 :** There are only 2 NCFs for *k* = 1 and those are *x*_1_ and 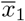. They are both *ChF*_0_ and *ChF*_1_. So, for *k* = 1 the three classes NCF, *ChF*_0_ and *ChF*_1_ are identical:

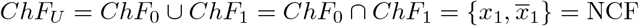
- ***k* = 2 :** There are 8 NCFs for *k* = 2. From the definitions of *ChF*_0_ (see Definition **4**^*∗*^) and *ChF*_1_ (see Definition **7**), the 6 *ChF*_0_s are 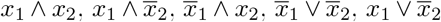 and 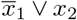 and the 6 *ChF*_1_s are 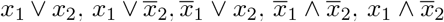 and 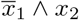. Hence we see that,

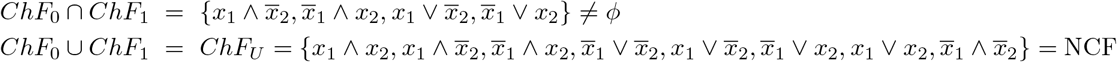 Hence, for *k* = 2, the classes *ChF*_0_ and *ChF*_1_ are not disjoint.
- ***k*** ≥ **3 :** Clearly, a multi-layer NCF can never simultaneously be a *ChF*_0_ and *ChF*_1_ since for each there is a strict dependency between the variable signs and the operators (∧ and ∨) within all but the last layer and no function can satisfy these distinct dependencies at once. So, let us consider a *k*-input (*k* ≥ 3) *ChF*_0_, *f* with single layer. Then, *m*_*last*_ = *k*. Now, *f* can have at most one inconsistency with the variable sign and the operator. Subsequently it follows that *f* will have at least (*k* − 1) inconsistencies between the variable signs and the operators required to get a *ChF*_1_. Since *k* ≥ 3, (*k* − 1) ≥ 2. This violates the sign-operator relationship for the *ChF*_1_ expression and hence *f* cannot be a *ChF*_1_. By symmetry, a *ChF*_1_ for *k* ≥ 3 cannot be a *ChF*_0_. Hence for *k* ≥ 3,

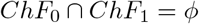

For all *k* ≥ 3 there exists multi-layer NCFs, some of which will always violate the condition of *ChF*_0_ (similarly *ChF*_1_) at least at one layer. Hence, for *k* ≥ 3,

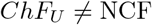

As an example, consider the 3-input NCF, *g* = *x*_1_ ∨ (*x*_2_ ∧ *x*_3_). Then, *g* violates the condition of being *ChF*_0_ in the first layer and the condition to be *ChF*_1_ in the second (last) layer and hence, this NCF is neither a *ChF*_0_ nor a *ChF*_1_.

### Property B.2.

The cardinality of *k*-input *ChF*_0_ or *ChF*_1_ with bias *P* are equal. As a result, the number of *k*-input *ChF*_0_ and *k*-input *ChF*_1_ is equal for any *k*.

*Proof*. Let us consider a map *f*_*p*_ whose domain is the class of *k*-input *ChF*_0_ with bias *P* and co-domain is *k*-input *ChF*_1_ with bias *P*. The map is defined by its flipping the signs of all the variables. Clearly, *f*_*p*_ is a one-to-one map. Hence, |*ChF*_0_|_*k,P*_ ≤ |*ChF*_1_|_*k,P*_ where, |*ChF*_0_|_*k,P*_ and |*ChF*_1_|_*k,P*_ correspond to the cardinality of the classes of *k*-input *ChF*_0_ and *ChF*_1_ with bias *P* respectively. In exactly similar manner one can define a one-to-one map from the class of *ChF*_1_ to the class of *ChF*_0_. Then, |*ChF*_1_|_*k,P*_ ≤ |*ChF*_0_|_*k,P*_. This proves that, |*ChF*_0_|_*k,P*_ = |*ChF*_1_|_*k,P*_. Hence, the total number of *k*-input *ChF*_0_ functions is equal to the total number of *k*-input *ChF*_1_ since,

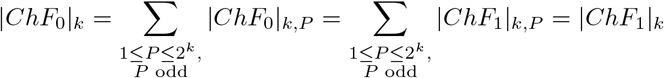

□

## Appendix C: The fraction of *ChF*_0_ (or *ChF*_1_) within NCF decreases exponentially with number of inputs

We provide here the derivation of the fraction of *k*-input *ChF*_0_ in NCFs for a given number of inputs *k*. From Eq. (6), the number of *k*-input NCFs is given by,

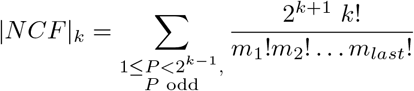

Again, from Eq. (11), the number of *k*-input *ChF*_0_ function is given by,

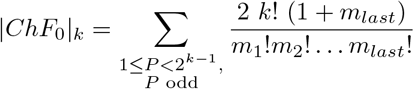

Hence, the fraction of *ChF*_0_ within NCF for some given value of *k* is,

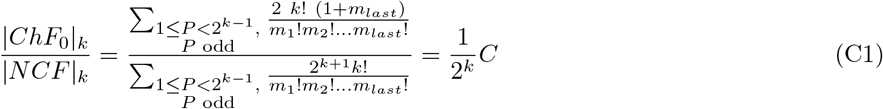

where,

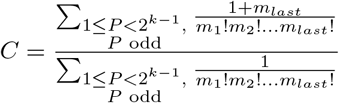

