## Supplementary material for "Preponderance of generalized chain functions in reconstructed Boolean models of biological networks": SI

### SUPPLEMENTARY INFORMATION
### for
### Preponderance of generalized chain functions in reconstructed Boolean models of biological networks

Suchetana Mitra,[1,2,*] Priyotosh Sil,[2,3,*] Ajay Subbaroyan,[2,3,*] Olivier C. Martin,[4,5,†] and Areejit Samal[2,3,†]

[1] *Indian Institute of Science Education and Research (IISER) Mohali, Punjab, India*
[2] *The Institute of Mathematical Sciences (IMSc), Chennai 600113, India*
[3] *Homi Bhabha National Institute (HBNI), Mumbai 400094, India*
[4] *Université Paris-Saclay, CNRS, INRAE, Univ Evry, Institute of Plant Sciences Paris-Saclay (IPS2), 91405 Orsay, France*
[5] *Université Paris-Cité, CNRS, INRAE, Institute of Plant Sciences Paris-Saclay (IPS2), 91405 Orsay, France*

#### SUPPLEMENTARY TABLES

TABLE S1. **Biological fixed points of the Pancreas cell differentiation model.** The 3 biological fixed points of the Pancreas cell differentiation model are the Exocrine, $\beta/\delta$ cell progenitor and $\alpha$/PP cell progenitor.

| **Cell types** | **Nodes** | | | | |
|---|---|---|---|---|---|
| | Pdx1 | Ptf1a | Ngn3 | Pax4 | Arx |
| Exocrine | 0 | 1 | 0 | 0 | 0 |
| $\beta/\delta$ cell progenitor | 1 | 0 | 1 | 1 | 0 |
| $\alpha$/PP cell progenitor | 0 | 0 | 1 | 0 | 1 |

TABLE S2. **Boolean functions of the published RSCN-2010 model in BoolNet format.** The column with header 'Target node name' contains the list of nodes whose regulation is captured by the corresponding row entry in the column 'Regulatory logic rule'. The symbols &, | and ! correspond to the logic operators AND, OR and NOT respectively.

| **Serial Number** | **Target node name** | **Regulatory logic rule** |
|---|---|---|
| 1 | SCR | ( SHR & SCR & !JKD & !MGP ) \| ( SHR & SCR & JKD & !MGP ) \| ( SHR & SCR & JKD & MGP ) |
| 2 | PLT | ARF |
| 3 | ARF | !AUXIAA |
| 4 | AUXIAA | !AUX |
| 5 | AUX | !AUX \| AUX |
| 6 | SHR | SHR |
| 7 | JKD | SHR & SCR |
| 8 | MGP | SHR & SCR & !WOX5 |
| 9 | WOX5 | (ARF & SHR & SCR & !MGP & !WOX5 ) \| (ARF & SHR & SCR & !MGP & WOX5 ) \| ( ARF & SHR & SCR & MGP & WOX5 ) |

* These authors contributed equally: Suchetana Mitra, Priyotosh Sil and Ajay Subbaroyan
† To whom correspondence should be addressed:


TABLE S3. **Biological fixed points recovered by the RSCN-2010 model.** The 4 biological fixed points recovered by the model are the Quiescent center (QC), Vascular initials (VI), Cortex-Endodermis initials (CEI) and Columella epidermis initials (CEpI).

| **Cell types** | **Nodes** | | | | | | | | |
|---|---|---|---|---|---|---|---|---|---|
| | SCR | PLT | ARF | AUXIAA | AUXIN | SHR | JKD | MGP | WOX5 |
| QC | 1 | 1 | 1 | 0 | 1 | 1 | 1 | 0 | 1 |
| VI | 0 | 1 | 1 | 0 | 1 | 1 | 0 | 0 | 0 |
| CEI | 1 | 1 | 1 | 0 | 1 | 1 | 1 | 1 | 0 |
| CEpI | 0 | 1 | 1 | 0 | 1 | 0 | 0 | 0 | 0 |

TABLE S4. **Boolean functions of the published RSCN-2020 model in BoolNet format.** The column with header 'Target node name' contains the list of nodes whose regulation is captured by the corresponding row entry in the column 'Regulatory logic rule'. The symbols &, | and ! correspond to the logic operators AND, OR and NOT respectively.

<table>
<tr><th>Serial Number</th><th>Target node name</th><th>Regulatory logic rule</th></tr>
<tr><td>1</td><td>CK</td><td>(PHB & !ARF) | !SHR</td></tr>
<tr><td>2</td><td>ARR1</td><td>!SCR & CK</td></tr>
<tr><td>3</td><td>SHY2</td><td>ARR1 & !AUX</td></tr>
<tr><td>4</td><td>AUXIAA</td><td>!AUX</td></tr>
<tr><td>5</td><td>ARF</td><td>!AUXIAA</td></tr>
<tr><td>6</td><td>ARF10</td><td>!(JKD & SHR) & !AUXIAA</td></tr>
<tr><td>7</td><td>ARF5</td><td>((PHB | PLT) & !(SHR & MGP)) & !SHY2 & !AUXIAA</td></tr>
<tr><td>8</td><td>XAL1</td><td>ARF</td></tr>
<tr><td>9</td><td>PLT</td><td>ARF5 | ARF | WOX5 | XAL1</td></tr>
<tr><td>10</td><td>AUX</td><td>AUX</td></tr>
<tr><td>11</td><td>SCR</td><td>SHR & JKD & SCR</td></tr>
<tr><td>12</td><td>SHR</td><td>SHR | (SCR & JKD)</td></tr>
<tr><td>13</td><td>MIR166</td><td>(SCR & SHR & !ARR1) | !PHB</td></tr>
<tr><td>14</td><td>PHB</td><td>((!ARR1 & PLT) | PHB) & !MIR166</td></tr>
<tr><td>15</td><td>JKD</td><td>!PHB & SHR & SCR</td></tr>
<tr><td>16</td><td>MGP</td><td>!ARF5 & SHR & SCR & MGP</td></tr>
<tr><td>17</td><td>WOX5</td><td>!ARF10 & ARF5 & !CLE40 & SCR & PLT</td></tr>
<tr><td>18</td><td>CLE40</td><td>!SHR</td></tr>
</table>

TABLE S5. **Biological fixed points recovered by the RSCN-2020 model.** The 6 biological fixed points recovered by the model are the Quiescent center (QC), Cortex/endodermis initial cell (CEI/ EndodermisPD), Peripheral Pro-vascular initials (P.ProvascularPD), Central Pro-vascular initials (C.ProvascularPD), Transition domain (C.ProvascularTD2) and Columella initials (Columella1).

| **Cell types** | **Nodes** | | | | | | | | | | | | | | | | | |
|---|---|---|---|---|---|---|---|---|---|---|---|---|---|---|---|---|---|---|
| | CK | ARR1 | SHY2 | AUXIAA | ARF | ARF10 | ARF5 | XAL1 | PLT | AUX | SCR | SHR | MIR166 | PHB | JKD | MGP | WOX5 | CLE40 |
| QC | 0 | 0 | 0 | 0 | 1 | 0 | 1 | 1 | 1 | 1 | 1 | 1 | 1 | 0 | 1 | 0 | 1 | 0 |
| CEI/EndodermisPD | 0 | 0 | 0 | 0 | 1 | 0 | 0 | 1 | 1 | 1 | 1 | 1 | 1 | 0 | 1 | 1 | 0 | 0 |
| P.ProvascularPD | 0 | 0 | 0 | 0 | 1 | 1 | 1 | 1 | 1 | 1 | 0 | 1 | 1 | 0 | 0 | 0 | 0 | 0 |
| C.ProvascularPD | 0 | 0 | 0 | 0 | 1 | 1 | 1 | 1 | 1 | 1 | 0 | 1 | 0 | 1 | 0 | 0 | 0 | 0 |
| C.ProvascularTD2 | 1 | 1 | 0 | 0 | 1 | 1 | 1 | 1 | 1 | 1 | 0 | 0 | 0 | 1 | 0 | 0 | 0 | 1 |
| Columella1 | 1 | 1 | 0 | 0 | 1 | 1 | 1 | 1 | 1 | 1 | 0 | 0 | 1 | 0 | 0 | 0 | 0 | 1 |

TABLE S6. **Numbers and fractions of $ChF_0$, $ChF_1$, $ChF_U$ and *non–*$ChF_U$ NCF in the BBM benchmark dataset.** $k$ is the number of inputs to a BF. The columns under the 'Number of BFs' grouping give the total number of BFs in the BBM benchmark dataset ('Total' column) and the numbers of BFs belonging to NCFs, $ChF_0$, $ChF_1$, $ChF_U$ and *non–*$ChF_U$ NCF types for various number of inputs. The columns under the 'Fraction of sub-types within NCFs' grouping give the fraction of the different sub-types of NCFs, namely, $ChF_0$, $ChF_1$, $ChF_U$ and *non–*$ChF_U$ NCF within the NCFs for various number of inputs.

| $k$ | Number of BFs | | | | | | Fraction of sub-types within NCFs | | | |
|---|---|---|---|---|---|---|---|---|---|---|
| | Total | NCF | $ChF_0$ | $ChF_1$ | $ChF_U$ | *non–*$ChF_U$ NCF | $ChF_0$ | $ChF_1$ | $ChF_U$ | *non–*$ChF_U$ NCF |
| 1 | 1998 | 1998 | 1998 | 1998 | 1998 | 0 | 1.0 | 1.0 | 1.0 | 0.0 |
| 2 | 1636 | 1625 | 1045 | 986 | 1625 | 0 | 0.64308 | 0.60677 | 1.0 | 0.0 |
| 3 | 915 | 886 | 279 | 467 | 746 | 140 | 0.3149 | 0.52709 | 0.84199 | 0.15801 |
| 4 | 558 | 502 | 97 | 218 | 315 | 187 | 0.19323 | 0.43426 | 0.62749 | 0.37251 |
| 5 | 366 | 297 | 31 | 152 | 183 | 114 | 0.10438 | 0.51178 | 0.61616 | 0.38384 |
| 6 | 224 | 158 | 25 | 64 | 89 | 69 | 0.15823 | 0.40506 | 0.56329 | 0.43671 |
| 7 | 119 | 80 | 7 | 22 | 29 | 51 | 0.0875 | 0.275 | 0.3625 | 0.6375 |
| 8 | 80 | 50 | 3 | 24 | 27 | 23 | 0.06 | 0.48 | 0.54 | 0.46 |
| 9 | 53 | 29 | 0 | 13 | 13 | 16 | 0.0 | 0.44828 | 0.44828 | 0.55172 |
| 10 | 41 | 16 | 0 | 9 | 9 | 7 | 0.0 | 0.5625 | 0.5625 | 0.4375 |

TABLE S7. **Relative enrichment ratios of $ChF_0$, $ChF_1$, $ChF_U$ and *non–*$ChF_U$ NCF within NCFs in the BBM benchmark dataset.** $k$ is the number of inputs to a BF. $ChF_0$, $ChF_1$, $ChF_U$ and *non–*$ChF_U$ NCF correspond to the chain-0, chain-1, generalized chain functions and NCFs that are not generalized chain functions respectively. The columns $ChF_0$, $ChF_1$, $ChF_U$ and *non–*$ChF_U$ NCF are the relative enrichment ratios (see Methods) for the associated sub-types within NCFs in the BBM benchmark dataset. $E_R > 1$ indicates that there is an enrichment of the sub-type within NCFs. NA implies that there are no BFs of the type *non–*$ChF_U$ NCF in theory.

| $k$ | Relative enrichment ratios ($E_R$) of sub-types within NCFs | | | |
|---|---|---|---|---|
| | $ChF_0$ | $ChF_1$ | $ChF_U$ | *non–*$ChF_U$ NCF |
| 1 | 1.0 | 1.0 | 1.0 | NA |
| 2 | 0.85744 | 0.80903 | 1.0 | NA |
| 3 | 0.77513 | 1.29745 | 1.03629 | 0.84274 |
| 4 | 0.9481 | 2.13078 | 1.53944 | 0.62882 |
| 5 | 1.02486 | 5.02514 | 3.025 | 0.48202 |
| 6 | 3.10739 | 7.95491 | 5.53115 | 0.48623 |
| 7 | 3.43675 | 10.80123 | 7.11899 | 0.6717 |
| 8 | 4.71326 | 37.70606 | 21.20966 | 0.47202 |
| 9 | 0.0 | 70.42801 | 35.21401 | 0.55884 |
| 10 | 0.0 | 176.74722 | 88.37361 | 0.4403 |

TABLE S8. **Statistical significance of the relative enrichments ($E_R$) of $ChF_0$, $ChF_1$, $ChF_U$ and *non*–$ChF_U$ NCF within NCFs for BFs in the BBM benchmark dataset.** $k$ is the number of inputs to a BF. The columns $ChF_0$, $ChF_1$, $ChF_U$ and *non*–$ChF_U$ NCF give the $p$-values associated with the relative enrichment (see Methods) of these sub-types of NCFs, within NCFs, for the BBM benchmark dataset. $p$-values $< 0.05$ are considered statistically significant. These $p$-values are used to assign the statistical significance stars in Fig. 1. Note that all 2-input NCFs are $ChF_U$s, hence it is meaningless to compute $p$-values for both $ChF_U$ and *non*–$ChF_U$ NCF.

| $k$ | $p$-values associated with $E_R$ for sub-types of NCFs | | | |
|---|---|---|---|---|
| | $ChF_0$ | $ChF_1$ | $ChF_U$ | *non*–$ChF_U$ NCF |
| 2 | 1 | 1 | NA | NA |
| 3 | 1.00000 | $1.63713 \times 10^{-13}$ | 0.00982 | 0.98756 |
| 4 | 0.70020 | $5.94764 \times 10^{-32}$ | $1.13295 \times 10^{-23}$ | 1 |
| 5 | 0.39584 | $3.61377 \times 10^{-71}$ | $1.28372 \times 10^{-54}$ | 1 |
| 6 | $1.33886 \times 10^{-07}$ | $1.44303 \times 10^{-41}$ | $1.90778 \times 10^{-47}$ | 1 |
| 7 | 0.00100 | $3.61870 \times 10^{-18}$ | $1.14534 \times 10^{-18}$ | 1 |
| 8 | 0.00379 | $3.88012 \times 10^{-34}$ | $1.18719 \times 10^{-31}$ | 1 |
| 9 | 0.16904 | $1.27064 \times 10^{-23}$ | $1.90290 \times 10^{-19}$ | 1 |
| 10 | 0.04972 | $8.38830 \times 10^{-22}$ | $8.44118 \times 10^{-19}$ | 1 |

TABLE S9. **Numbers and fractions of $ChF_0$, $ChF_1$, $ChF_U$ and *non*–$ChF_U$ NCF in the MCBF dataset.** $k$ is the number of inputs to a BF. The columns under the 'Number of BFs' grouping give the total number of BFs in the MCBF dataset ('Total' column) and the numbers of BFs belonging to NCFs, $ChF_0$, $ChF_1$, $ChF_U$ and *non*–$ChF_U$ NCF types for various number of inputs. The columns under the 'Fraction of sub-types within NCFs' grouping give the fraction of the different sub-types of NCFs, namely, $ChF_0$, $ChF_1$, $ChF_U$ and *non*–$ChF_U$ NCF within the NCFs for various number of inputs.

| $k$ | Number of BFs | | | | | | Fraction of sub-types within NCFs | | | |
|---|---|---|---|---|---|---|---|---|---|---|
| | Total | NCF | $ChF_0$ | $ChF_1$ | $ChF_U$ | *non*–$ChF_U$ NCF | $ChF_0$ | $ChF_1$ | $ChF_U$ | *non*–$ChF_U$ NCF |
| 1 | 934 | 934 | 934 | 934 | 934 | 0 | 1.0 | 1.0 | 1.0 | 0.0 |
| 2 | 687 | 671 | 392 | 503 | 671 | 0 | 0.5842 | 0.74963 | 1.0 | 0.0 |
| 3 | 412 | 378 | 116 | 199 | 315 | 63 | 0.30688 | 0.52646 | 0.83333 | 0.16667 |
| 4 | 258 | 230 | 31 | 124 | 155 | 75 | 0.13478 | 0.53913 | 0.67391 | 0.32609 |
| 5 | 156 | 120 | 10 | 63 | 73 | 47 | 0.08333 | 0.525 | 0.60833 | 0.39167 |
| 6 | 107 | 67 | 10 | 26 | 36 | 31 | 0.14925 | 0.38806 | 0.53731 | 0.46269 |
| 7 | 51 | 34 | 4 | 10 | 14 | 20 | 0.11765 | 0.29412 | 0.41176 | 0.58824 |
| 8 | 45 | 27 | 2 | 9 | 11 | 16 | 0.07407 | 0.33333 | 0.40741 | 0.59259 |
| 9 | 19 | 7 | 0 | 2 | 2 | 5 | 0.0 | 0.28571 | 0.28571 | 0.71429 |
| 10 | 13 | 3 | 0 | 1 | 1 | 2 | 0.0 | 0.33333 | 0.33333 | 0.66667 |

TABLE S10. **Relative enrichment ratios of $ChF_0$, $ChF_1$, $ChF_U$ and *non–*$ChF_U$ NCF within NCFs in the MCBF dataset** $k$ is the number of inputs to a BF. $ChF_0$, $ChF_1$, $ChF_U$ and *non–*$ChF_U$ NCF correspond to the chain-0, chain-1, generalized chain functions and NCFs that are not generalized chain functions respectively. The columns $ChF_0$, $ChF_1$, $ChF_U$ and *non–*$ChF_U$ NCF are the relative enrichment ratios (see Methods) for the associated sub-types within NCFs in the MCBF dataset. $E_R > 1$ indicates that there is an enrichment of the sub-type within NCFs. NA implies that there are no BFs of the type *non–*$ChF_U$ NCF in theory.

| $k$ | Relative enrichment ratios ($E_R$) of sub-types within NCFs | | | |
|---|---|---|---|---|
| | $ChF_0$ | $ChF_1$ | $ChF_U$ | *non–*$ChF_U$ NCF |
| 1 | 1.0 | 1.0 | 1.0 | NA |
| 2 | 0.77894 | 0.9995 | 1.0 | NA |
| 3 | 0.75539 | 1.29589 | 1.02564 | 0.88889 |
| 4 | 0.66133 | 2.64533 | 1.65333 | 0.55046 |
| 5 | 0.81824 | 5.1549 | 2.98657 | 0.49185 |
| 6 | 2.93115 | 7.62099 | 5.27607 | 0.51515 |
| 7 | 4.62085 | 11.55211 | 8.08648 | 0.6198 |
| 8 | 5.81884 | 26.18477 | 16.0018 | 0.60807 |
| 9 | 0.0 | 44.88818 | 22.44409 | 0.7235 |
| 10 | 0.0 | 104.7391 | 52.36955 | 0.67094 |

TABLE S11. **Statistical significance of the relative enrichments ($E_R$) of $ChF_0$, $ChF_1$, $ChF_U$ and *non–*$ChF_U$ NCF within NCFs for BFs in the MCBF dataset.** $k$ is the number of inputs to a BF. The columns $ChF_0$, $ChF_1$, $ChF_U$ and *non–*$ChF_U$ NCF give the $p$-values associated with the relative enrichment (see Methods) of these sub-types of NCFs, within NCFs, for the MCBF dataset. $p$-values $< 0.05$ are considered statistically significant. These $p$-values are used to assign the statistical significance stars in Fig. S3. Note that all 2-input NCFs are $ChF_U$s, hence it is meaningless to compute $p$-values for both $ChF_U$ and *non–*$ChF_U$ NCF.

| $k$ | $p$-values associated with $E_R$ for sub-types of NCFs | | | |
|---|---|---|---|---|
| | $ChF_0$ | $ChF_1$ | $ChF_U$ | *non–*$ChF_U$ NCF |
| 2 | 1 | 0.49407518307496 | NA | NA |
| 3 | 0.99996 | $9.53875 \times 10^{-07}$ | 0.13412 | 0.83431 |
| 4 | 0.99561 | $8.77878 \times 10^{-30}$ | $1.08696 \times 10^{-16}$ | 1 |
| 5 | 0.68767 | $6.46209 \times 10^{-32}$ | $8.96155 \times 10^{-23}$ | 1 |
| 6 | 0.00054 | $6.69661 \times 10^{-18}$ | $8.58220 \times 10^{-20}$ | 1 |
| 7 | 0.00161 | $4.84767 \times 10^{-10}$ | $2.94472 \times 10^{-11}$ | 1.00000 |
| 8 | 0.00480 | $7.73712 \times 10^{-13}$ | $9.03023 \times 10^{-13}$ | 1.00000 |
| 9 | 0.04371 | $8.85440 \times 10^{-06}$ | $6.94878 \times 10^{-05}$ | 0.99674 |
| 10 | 0.00952 | $3.03207 \times 10^{-05}$ | 0.00012 | 0.98103 |

TABLE S12. **Numbers and fractions of $ChF_0$, $ChF_1$, $ChF_U$ and *non–*$ChF_U$ NCF in the Harris dataset.** $k$ is the number of inputs to a BF. The columns under the 'Number of BFs' grouping give the total number of BFs in the Harris dataset ('Total' column) and the numbers of BFs belonging to NCFs, $ChF_0$, $ChF_1$, $ChF_U$ and *non–*$ChF_U$ NCF types for various number of inputs. The columns under the 'Fraction of sub-types within NCFs' grouping give the fraction of the different sub-types of NCFs, namely, $ChF_0$, $ChF_1$, $ChF_U$ and *non–*$ChF_U$ NCF within the NCFs for various number of inputs.

| $k$ | Number of BFs | | | | | | Fraction of sub-types within NCFs | | | |
|---|---|---|---|---|---|---|---|---|---|---|
| | Total | NCF | $ChF_0$ | $ChF_1$ | $ChF_U$ | *non–*$ChF_U$ NCF | $ChF_0$ | $ChF_1$ | $ChF_U$ | *non–*$ChF_U$ NCF |
| 1 | 2 | 2 | 2 | 2 | 2 | 0 | 1.0 | 1.0 | 1.0 | 0.0 |
| 2 | 9 | 9 | 9 | 3 | 9 | 0 | 1.0 | 0.33333 | 1.0 | 0.0 |
| 3 | 71 | 71 | 59 | 3 | 62 | 9 | 0.83099 | 0.04225 | 0.87324 | 0.12676 |
| 4 | 38 | 35 | 26 | 0 | 26 | 9 | 0.74286 | 0.0 | 0.74286 | 0.25714 |
| 5 | 19 | 16 | 11 | 0 | 11 | 5 | 0.6875 | 0.0 | 0.6875 | 0.3125 |

TABLE S13. **Relative enrichment ratios of $ChF_0$, $ChF_1$, $ChF_U$ and *non–*$ChF_U$ NCF within NCFs in the Harris dataset** $k$ is the number of inputs to a BF. $ChF_0$, $ChF_1$, $ChF_U$ and *non–*$ChF_U$ NCF correspond to the chain-0, chain-1, generalized chain functions and NCFs that are not generalized chain functions respectively. The columns $ChF_0$, $ChF_1$, $ChF_U$ and *non–*$ChF_U$ NCF are the relative enrichment ratios (see Methods) for the associated sub-types within NCFs in the Harris dataset. $E_R > 1$ indicates that there is an enrichment of the sub-type within NCFs. NA implies that there are no BFs of the type *non–*$ChF_U$ NCF in theory.

| $k$ | Relative enrichment ratios ($E_R$) of sub-types within NCFs | | | |
|---|---|---|---|---|
| | $ChF_0$ | $ChF_1$ | $ChF_U$ | *non–*$ChF_U$ NCF |
| 1 | 1.0 | 1.0 | 1.0 | NA |
| 2 | 1.33333 | 0.44444 | 1.0 | NA |
| 3 | 2.0455 | 0.10401 | 1.07476 | 0.67606 |
| 4 | 3.64495 | 0.0 | 1.82248 | 0.43408 |
| 5 | 6.75046 | 0.0 | 3.37523 | 0.39243 |

TABLE S14. **Statistical significance of the relative enrichments ($E_R$) of $ChF_0$, $ChF_1$, $ChF_U$ and *non–*$ChF_U$ NCF within NCFs for BFs in the Harris dataset.** $k$ is the number of inputs to a BF. The columns $ChF_0$, $ChF_1$, $ChF_U$ and *non–*$ChF_U$ NCF give the $p$-values associated with the relative enrichment (see Methods) of these sub-types of NCFs, within NCFs, for the Harris dataset. $p$-values $< 0.05$ are considered statistically significant. These $p$-values are used to assign the statistical significance stars in Fig. S3. Note that all 2-input NCFs are $ChF_U$s, hence it is meaningless to compute $p$-values for both $ChF_U$ and *non–*$ChF_U$ NCF.

| $k$ | $p$-values associated with $E_R$ for sub-types of NCFs | | | |
|---|---|---|---|---|
| | $ChF_0$ | $ChF_1$ | $ChF_U$ | *non–*$ChF_U$ NCF |
| 2 | 0 | 0.99001 | NA | NA |
| 3 | $3.17607 \times 10^{-14}$ | 1.00000 | 0.06557 | 0.87972 |
| 4 | $9.14592 \times 10^{-13}$ | 0.99966 | $1.32004 \times 10^{-05}$ | 0.99994 |
| 5 | $1.52758 \times 10^{-09}$ | 0.82068 | $4.04322 \times 10^{-06}$ | 0.99996 |

TABLE S15. **Nodewise enumeration of the number of BFs that satisfy biological constraints for the Pancreas cell differentiation GRN.** 'Nodes' are the names of the nodes in the network and $k$ is the associated number of inputs to that node. The columns $ChF_0$, $ChF_1$ and $ChF_U$ give the number of chain-0, chain-1 and generalized chain functions that satisfy biological fixed point constraints and sign conforming constraints at each node of the Pancreas cell differentiation GRN. Imposing $ChF_U$ leads to 3600 Boolean models.

| Nodes | Inputs | $ChF_0$ | $ChF_1$ | $ChF_U$ |
|---|---|---|---|---|
| Pdx1 | 1 | 1 | 1 | 1 |
| Ptf1a | 3 | 4 | 6 | 10 |
| Ngn3 | 3 | 4 | 6 | 10 |
| Pax4 | 3 | 4 | 2 | 6 |
| Arx | 3 | 4 | 2 | 6 |

TABLE S16. **Nodewise enumeration of the number of BFs that satisfy biological constraints for the RSCN-2020 GRN.** 'Nodes' are the names of the nodes in the network and $k$ is the associated number of inputs to that node. The columns $ChF_0$, $ChF_1$ and $ChF_U$ columns give the number of chain-0, chain-1 and generalized chain functions that satisfy biological fixed point constraints and sign conforming constraints at each node of the RSCN-2020 GRN. Note here that for the ARF10 node, there is no generalized chain function that satisfies the above-mentioned constraints. Thus for ARF10, the allowed BF was restricted to NCF leading to 4 BFs for that node. Imposing $ChF_U$ on all nodes except for ARF10 (for which NCF is imposed) leads to a total of 645120 models.

| Nodes | Inputs | $ChF_0$ | $ChF_1$ | $ChF_U$ |
|---|---|---|---|---|
| CK | 3 | 1 | 0 | 1 |
| ARR1 | 2 | 1 | 1 | 1 |
| SHY2 | 2 | 1 | 1 | 1 |
| AUXIAA | 1 | 1 | 1 | 1 |
| ARF | 1 | 1 | 1 | 1 |
| ARF10 | 3 | 0 | 0 | 0 |
| ARF5 | 6 | 16 | 32 | 48 |
| XAL1 | 1 | 1 | 1 | 1 |
| PLT | 4 | 0 | 1 | 1 |
| AUX | 1 | 1 | 1 | 1 |
| SCR | 3 | 1 | 0 | 1 |
| SHR | 3 | 0 | 1 | 1 |
| MIR166 | 4 | 2 | 0 | 2 |
| PHB | 4 | 7 | 5 | 12 |
| JKD | 3 | 2 | 0 | 2 |
| MGP | 4 | 8 | 2 | 10 |
| WOX5 | 5 | 7 | 0 | 7 |
| CLE40 | 1 | 1 | 1 | 1 |

#### SUPPLEMENTARY FIGURES

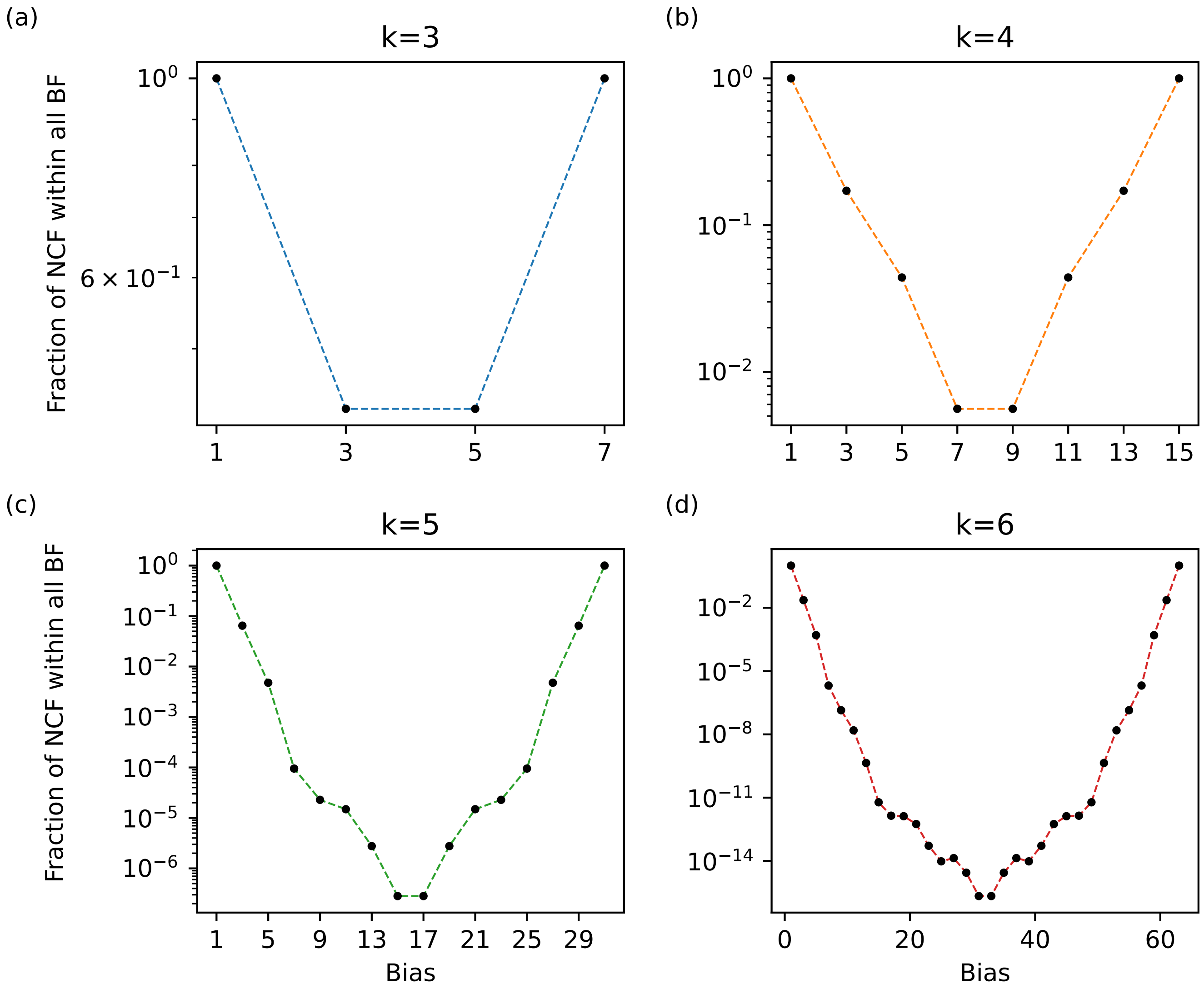


FIG. S1. **Bias-wise fraction of NCFs within all BFs.** For a given number of inputs ($k$), the fraction of NCFs in all BFs ($y$-axis) is plotted as a function of the bias ($1 \leq P \leq 2^k - 1$ for odd $P$) ($x$-axis). Subplots correspond to different number of inputs $k = 3, 4, 5$ and 6.

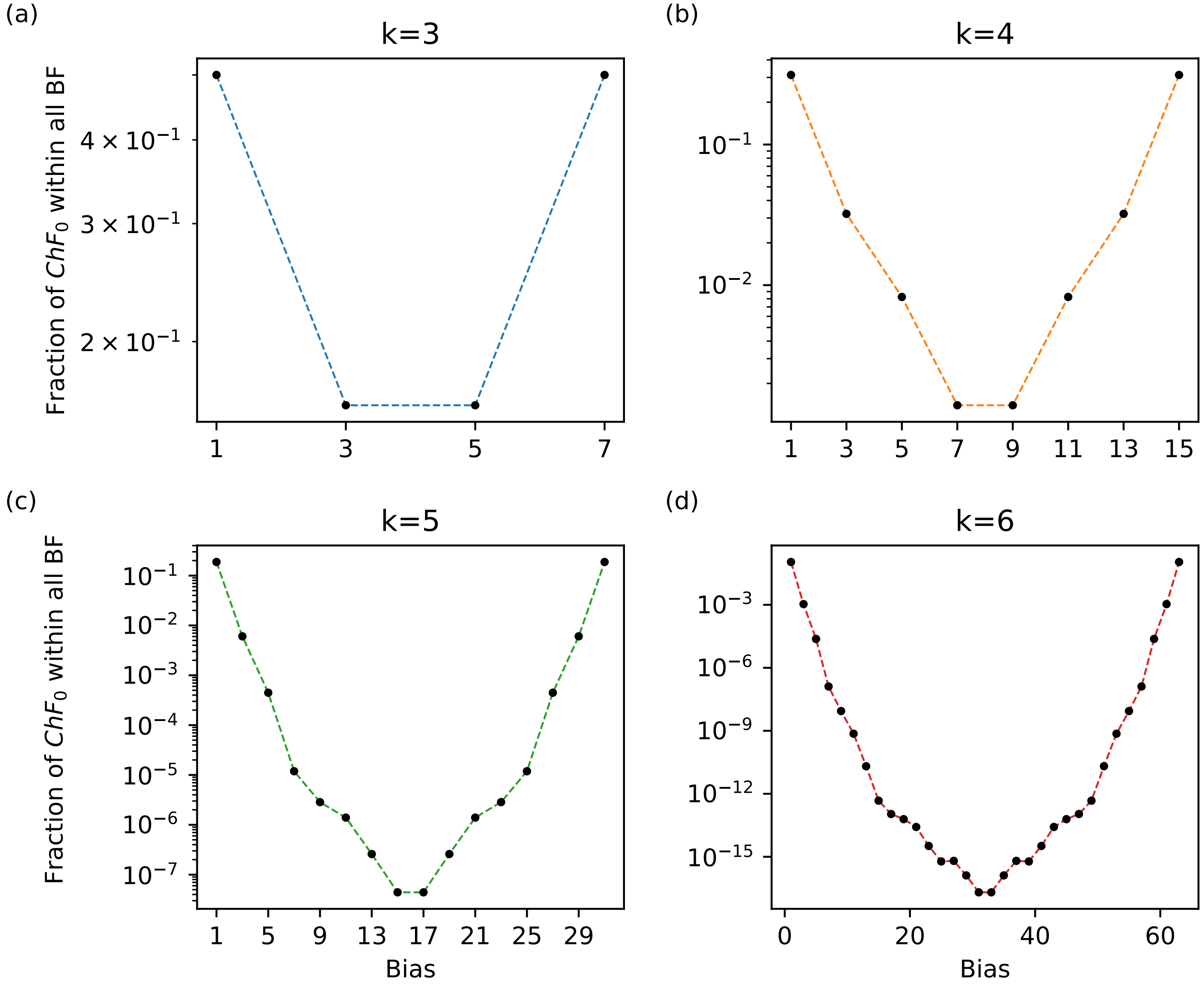


FIG. S2. **Bias-wise fraction of $ChF_0$ (or $ChF_1$) within all BFs.** For a given number of inputs ($k$), the fraction of $ChF_0$ (or $ChF_1$) in all BFs ($y$-axis) is plotted as a function of the bias ($1 \le P \le 2^k - 1$ for odd $P$) ($x$-axis). Subplots correspond to different number of inputs $k = 3, 4, 5$ and 6.

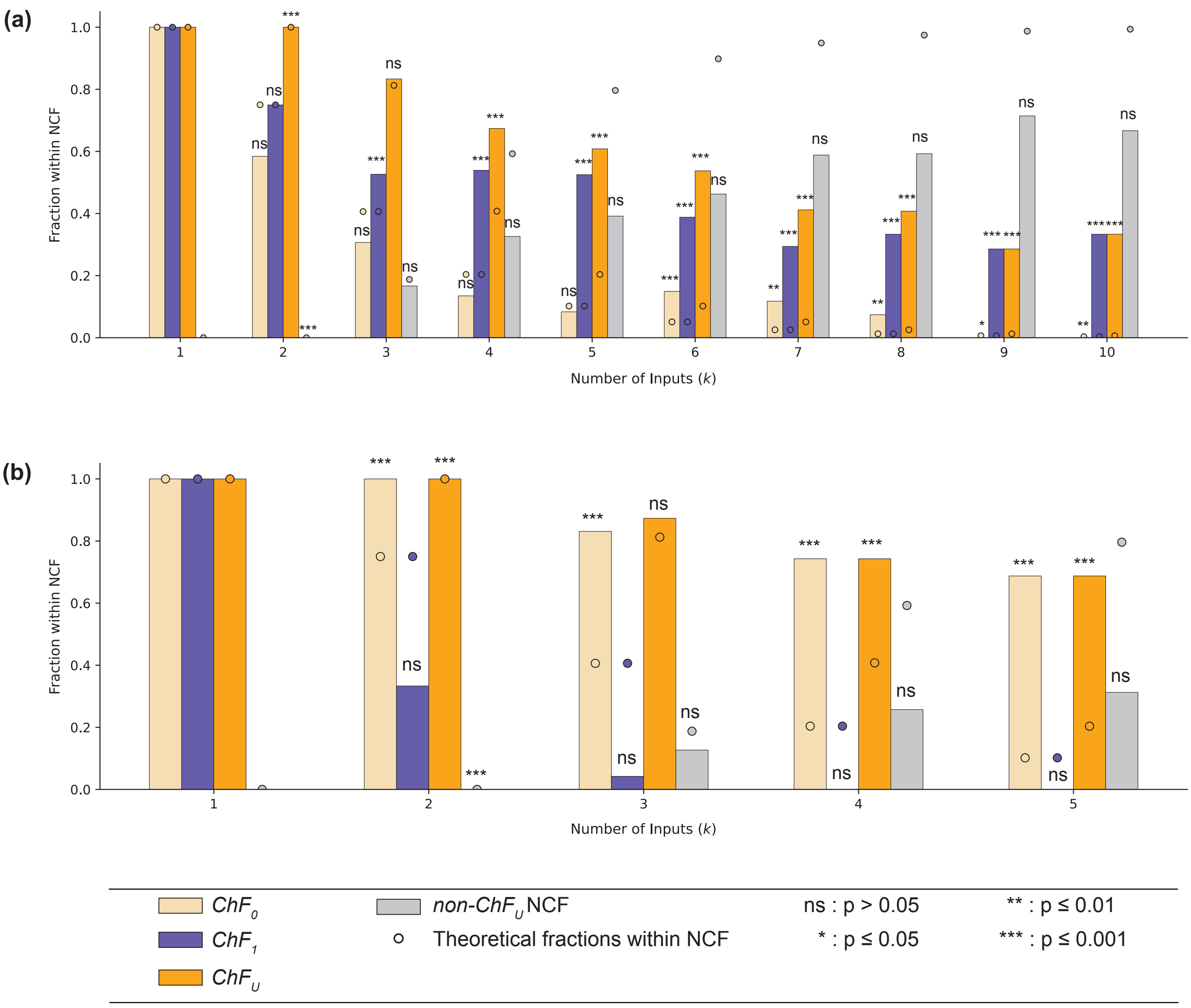


FIG. S3. **Fractions of $ChF_0$, $ChF_1$, $ChF_U$ and $non-ChF_U NCF$ within NCFs in the MCBF dataset and Harris dataset (a)** This sub-figure shows the fraction of $ChF_0$, $ChF_1$, $ChF_U$ and $non-ChF_U NCF$ within NCFs, in theory and in the MCBF dataset as dots and colored bars respectively. The relative enrichments of $ChF_1$s and $ChF_U$s within NCF are statistically significant for $k \geq 3$ and $k \geq 2$ (except for $k = 3$) respectively. **(b)** This sub-figure shows the fractions of $ChF_0$, $ChF_1$, $ChF_U$ and $non-ChF_U NCF$ within NCF in theory and in the Harris dataset as dots and colored bars respectively. The relative enrichments of $ChF_0$s and $ChF_U$s within NCFs are statistically significant for $k \geq 2$ (except for $ChF_U$ at $k = 3$).
